# Accurate, scalable, and fully automated inference of species trees from raw genome assemblies using ROADIES

**DOI:** 10.1101/2024.05.27.596098

**Authors:** Anshu Gupta, Siavash Mirarab, Yatish Turakhia

## Abstract

Inference of species trees plays a crucial role in advancing our understanding of evolutionary relationships and has immense significance for diverse biological and medical applications. Extensive genome sequencing efforts are currently in progress across a broad spectrum of life forms, holding the potential to unravel the intricate branching patterns within the tree of life. However, estimating species trees starting from raw genome sequences is quite challenging, and the current cutting-edge methodologies require a series of error-prone steps that are neither entirely automated nor standardized. In this paper, we present ROADIES, a novel pipeline for species tree inference from raw genome assemblies that is fully automated, easy to use, scalable, free from reference bias, and provides flexibility to adjust the tradeoff between accuracy and runtime. The ROADIES pipeline eliminates the need to align whole genomes, choose a single reference species, or pre-select loci such as functional genes found using cumbersome annotation steps. Moreover, it leverages recent advances in phylogenetic inference to allow multi-copy genes, eliminating the need to detect orthology. Using the genomic datasets released from large-scale sequencing consortia across three diverse life forms (placental mammals, pomace flies, and birds), we show that ROADIES infers species trees that are comparable in quality with the state-of-the-art approaches but in a fraction of the time. By incorporating optimal approaches and automating all steps from assembled genomes to species and gene trees, ROADIES is poised to improve the accuracy, scalability, and reproducibility of phylogenomic analyses.

**Code and Data availability:** The source code of ROADIES is freely available under the MIT License on GitHub (https://github.com/TurakhiaLab/ROADIES), and the documentation for ROADIES is available at https://turakhia.ucsd.edu/ROADIES/. The details of the input datasets used in the manuscript are listed in Supplementary Tables 1-3. All inferred gene trees and species trees are to be deposited to Dryad with links to be made available on the aforementioned GitHub repository.

## Introduction

The rapid progress in genome sequencing technologies and assembly methods has ushered in an era wherein accurate and complete genome assemblies of diverse species are being produced at an unprecedented rate. For example, tens of thousands to millions of eukaryotic species are expected to be sequenced and assembled in the next decade through various ongoing projects ^1–3^. These genomes hold the promise to resolve long-standing questions surrounding the evolutionary relationship of species (species trees) and illuminate differences in evolutionary history across the genome (gene trees) ^4–6^. These *phylogenomic* analyses, however, are cumbersome and involve many intricate steps. Thus, the rapid genomic data expansion is creating a pressing need to automate phylogenomic inference pipelines that follow best practices to infer accurate gene trees and species starting from raw genome assemblies ^7^.

Despite this need, automation of accurate species tree inference from genome assemblies has been an enduring challenge. While there is no universally agreed-upon method for accurately and reliably inferring species trees, modern phylogenetic pipelines are increasingly adopting methods that account for gene tree discordance and are statistically consistent under various models of genome evolution ^8–11^. However, these pipelines are notoriously hard to automate and involve many error-prone steps. For example, a key challenge is that the current practice generally relies on precise gene annotations and orthology inference ^4,12,13^, which is an art form in itself, demanding substantial human involvement and specialized knowledge. The complex and nuanced steps involved in the gene tree inference process include i) selecting and annotating loci (which are often but not always functional genes) in exemplar species, ii) identifying orthologous regions corresponding to those genes in other species, and iii) determining the most suitable method and the set of parameters to align and infer the evolutionary trees of the genes ^14–22^. The first two steps could rely on gene annotation or whole genome alignment, each of which is a difficult procedure. Due to these reasons, full automation of species tree inference has rarely been attempted. While there have been efforts to automate parts of the phylogenetic workflow ^23^, the only fully automated tools for species tree inference that currently exist focus on simpler methods that involve fewer parameters and algorithmic stages, such as distance-based methods ^24^, which have considerable limitations in terms of the accuracy, reliability, and interpretability ^25^.

In this paper, we attempt to automate accurate, reliable, fast, and scalable inference of species trees starting from raw genome assemblies. Our automated tool is called ROADIES, which is an abbreviation for “*<u>R</u>eference-free, <u>O</u>rthology-free, <u>A</u>lignment-free, <u>Di</u>scordance-aware <u>E</u>stimation of <u>S</u>pecies Trees*,” as it includes the following key features:

1. **Reference-free:** ROADIES does not require a reference species, gene, set, or genome annotations. It also does not depend on a single genome as a reference for alignments or gene selection and is, therefore, free from reference bias.
2. **Orthology-free:** ROADIES allows multi-copy gene trees (inferred from homologous regions) and does not require the challenging and error-prone step of orthologous detection prior to gene tree inference. Note that we use the term “gene” to refer to c-genes (i.e., coalescent genes: short regions of the genome that are ideally recombination-free) ^26,27^, not the more common usage meaning functional genes.
3. **Alignment-free:** ROADIES does not require any input alignment to be provided, as all alignments required for gene tree inference are constructed within the ROADIES pipeline itself.
4. **Discordance-aware:** ROADIES uses a state-of-the-art and statistically consistent discordance-aware method to combine gene trees into a species tree.

ROADIES has been evaluated for large-scale phylogenetic inference on three diverse datasets – mammals, birds, and pomace flies. We found that the phylogenetic trees generated by ROADIES were largely concordant with the findings of careful, large-scale studies on these datasets that employed state-of-the-art practices and discordance-aware methods for deducing phylogenetic relationships ^12,28–30^. A notable aspect of ROADIES is the random sampling of c-genes across the genome, which is in line with the theory underlying phylogenomic inference ^10,31^, suggestions from empiricists ^27,32^, and recent phylogenomic analyses ^6,33^. ROADIES is highly parallelized, scalable, and configurable. We estimate that ROADIES requires a fraction of the runtime and no human intervention compared to the state-of-the-art approaches that include either gene annotation or multiple whole genome alignment (m-WGA) ^34^ and orthology inference. ROADIES inferences are far more accurate than a currently available fully-automated approach, MashTree ^24^.

We expect ROADIES to find broad applications. These range from constructing guide trees for whole-genome alignments ^34^ to contributing to various evolutionary biology studies and aiding in the resolution of the Tree of Life ^35^. To accommodate a wide array of users with varying requirements and constraints, ROADIES offers three distinct operational modes: ‘*accurate*,’ ‘*balanced*,’ and ‘*fast*.’ Each mode presents a different trade-off between runtime and accuracy, enabling users to select the option that best suits their needs.

## Results

### ROADIES Overview

ROADIES is a fully automated pipeline to produce species and gene trees from raw genome assemblies (Fig. 1A). There are two ways in which ROADIES differs from conventional discordance-aware coalescent pipelines for species tree inference (Fig. 1B). *First*, most conventional pipelines are designed only to handle single-copy gene trees reconstructed from sets of orthologous genes. However, separating orthologs from paralogs is challenging, requiring domain expertise in some cases ^9,19,20,22,36^. ROADIES eliminates the need for orthology inference by leveraging ASTRAL-Pro2 ^37^, a recently developed discordance-aware summary method that can produce accurate species trees from multi-copy gene trees, i.e., gene trees in which all homologs are represented without needing to separate orthologs from paralogs. In effect, ROADIES leaves teasing out orthology and paralogy based on the inferred gene family trees to ASTRAL-Pro2. *Second*, most conventional pipelines rely on gene annotation, typically of protein-coding genes, or use m-WGA to select evenly sampled loci from single-copy regions ^6,33,38^ (Fig. 1B). Annotation tends to be computationally expensive and requires significant domain expertise^39^. Besides, protein-coding genes have been criticized as input to phylogenomics due to their widespread deviations from sequence evolution models ^4,6,40,41^. The use of m-WGA is also complicated by the difficulty of inferring such alignments, which itself requires a guide tree ^34^ and finding single-copy regions. Instead of relying on protein-coding gene annotations or m-WGA, ROADIES samples random sequences from different input genomes. ROADIES’ gene sampling strategy has multiple benefits: (i) it eliminates the need to perform costly annotations of protein-coding genes or other regions on input genomes, (ii) it eliminates reference bias, as instead of a single genome, genes are sampled randomly from all input genomes with a uniform distribution, (iii) by not restricting the genes from coding regions of the genome, this strategy allows intergenic regions (expected to follow assumptions of neutral evolution better) to be selected, and (iv) it does not require costly m-WGA and a starting guide tree. The aforementioned differences with respect to conventional methods greatly simplify the computational process in ROADIES, specifically for automation, and help ROADIES achieve orders of magnitude gain in performance (Fig. 1D) without sacrificing accuracy (Fig. 1E).

**Figure 1:**
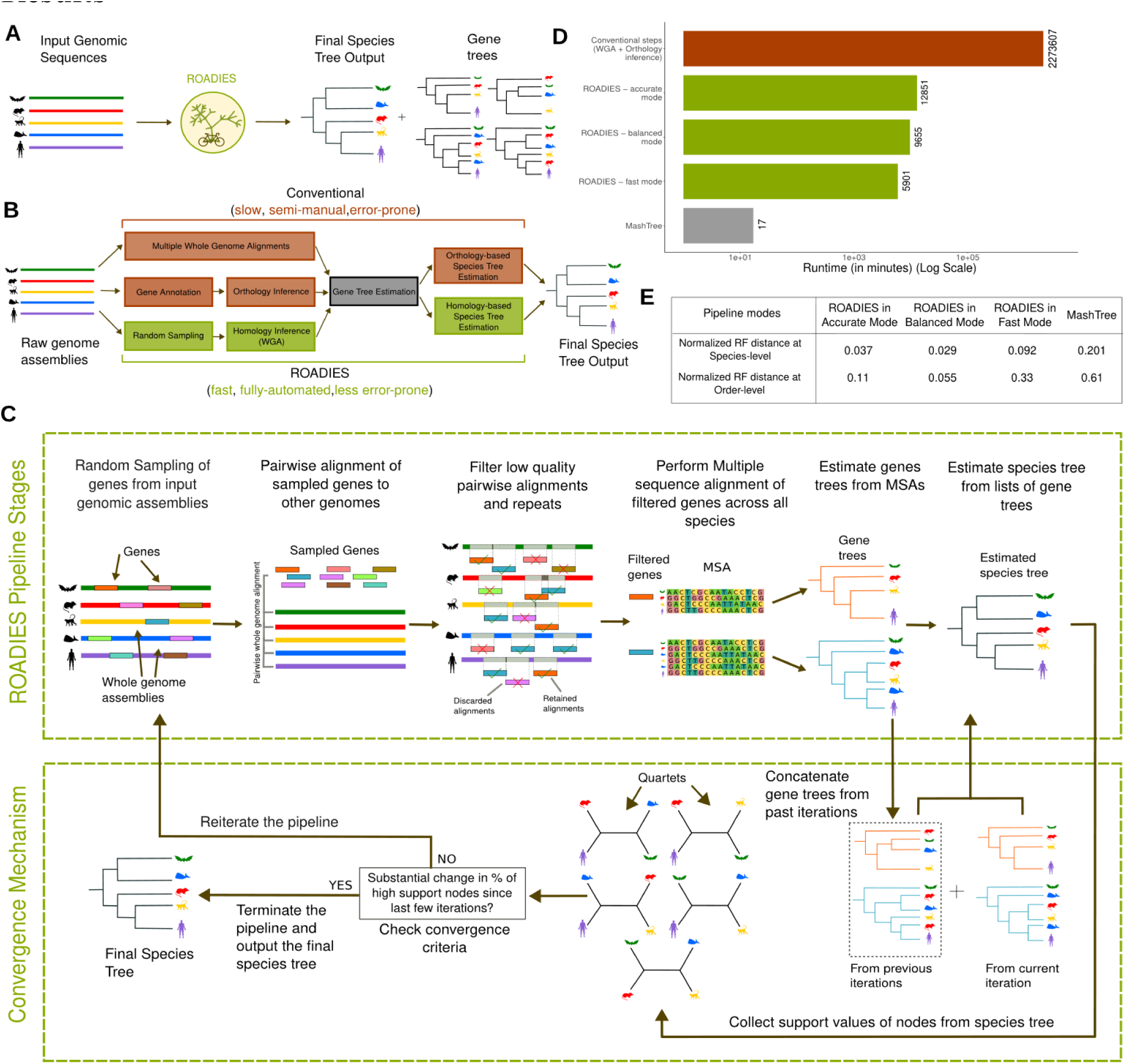
An overview of the ROADIES pipeline. **(A)** ROADIES input and output (**B**) A comparison of the different stages involved in the species tree inference in the conventional approaches and ROADIES. (**C**) A detailed view of the ROADIES pipeline’s various stages and the convergence mechanism. (**D**) Runtime estimates for inferring the species tree of 240 placental mammals from raw genome assemblies using the conventional approaches, the three modes (accurate, balanced, and fast) of ROADIES and MashTree ^24^. For conventional approaches, we only include the runtime estimates of the whole-genome alignments and orthology estimation stages. All runtime projections are provided for a 16-core AWS EC2 instance (r6a.4xlarge) (Methods). (**E**) Normalized Robinson-Foulds distance (normRF, lower is better) between the reference tree of 240 placental mammals from the Zoonomia consortium ^12^ and the species trees inferred by the different modes of operation of ROADIES and MashTree.

ROADIES is an iterative approach (Fig. 1C) and provides a high degree of configurability in parameters; it includes three convenient modes of operation: accurate, balanced, and fast (Methods). All modes keep iterating till a confident and stable tree is achieved. In the first iteration, ROADIES starts with a default of 250 randomly sampled genes (GENE_COUNT = 500), each of 500bp (LENGTH = 500), selected from distinct randomly sampled input genomes. It then finds homologous regions corresponding to these genes in all remaining genomes using LASTZ ^42^. Next, in accurate mode, for each gene, a multiple-sequence alignment (MSA) of all its homologs in all genomes is inferred using PASTA ^43^, followed by multi-copy gene tree inference using RaxML-NG ^44^. In balanced mode, the multi-copy gene tree inference is done using FastTree ^45^, an approximate likelihood technique that is a lot faster than RaxML-NG ^44^. In fast mode, the MSA step is eliminated entirely by producing gene trees using MashTree ^24^, a neighbor-joining-based multi-copy gene tree of Mash distances inferred from homologs. In all modes, ASTRAL-Pro2 ^37^ is used at the end to combine the multi-copy gene trees into a single species tree. ASTRAL-Pro2 ^37^ also reports confidence scores for each branch in the form of local posterior probability (localPP)^46^. ROADIES begins a new iteration with the doubling of gene count if a stopping criterion is not met; by default, we test if the number of highly-confident branches (i.e., localPP >= 0.95) changes by less than 1% from the previous iteration. Appropriate filtering strategies are used between stages to ensure the quality of final results (see Methods for full details).

### ROADIES estimates an accurate phylogenetic tree of 240 placental mammals

We evaluated the accuracy and performance of ROADIES in the accurate mode using the genome assemblies of 240 placental mammals provided by the Zoonomia consortium ^12^ (Supplementary Table 1, Methods). The assemblies include representative species from all 20 orders of the placental mammalian phylogeny (Supplementary Table 1). For accuracy comparison, we used the tree topology provided by Zoonomia ^12^ as a representative of the state-of-the-art scientific literature. We found that the ROADIES-inferred phylogeny is largely in agreement with the Zoonomia phylogeny (Fig. 2). At the species level, the normalized Robinson-Foulds (normRF) distance ^47^ between the two phylogenies is only 0.037. In both order-level phylogenetic trees, every species is accurately assigned to its respective phylogenetic order. Minor differences are noticeable in the arrangement of the orders (Fig. 2B). At the order level, only two relationships changed (nodes 1 and 2, Fig. 2B), resulting in a normRF distance of 0.11. These contested relationships are resolved with low confidence (localPP < 0.78) compared to other highly confident branches (localPP >= 0.95). ROADIES stochastically picks the number of genes from each species as a reference and finds an almost uniform representation of each species among the gene trees. However, some species, such as rodents, have comparatively fewer aligned genes (Fig. 2A) likely because of its significantly faster evolution rate compared to other neighboring orders ^48,49^, which makes it challenging to infer homology. We next compare the two phylogenies (reference and ROADIES) with a specific focus on historically contested branches:

**Figure 2:**
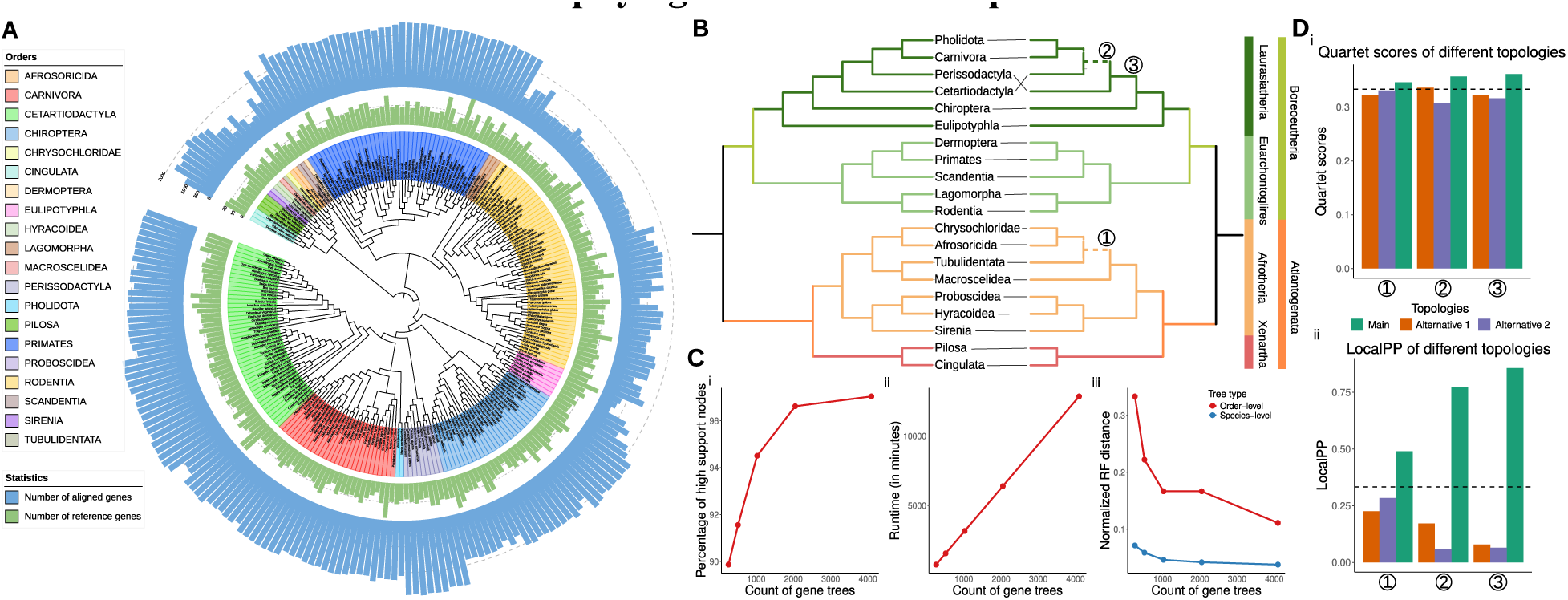
ROADIES results evaluated on the dataset of 240 placental mammals (in the accurate mode of operation). **(A)** The species-level phylogenetic tree of 240 placental mammals estimated by ROADIES. The number of genes aligned to each species (blue) and the count of genes sampled from each species (green) ^12^ are also shown. **(B)** Order-level trees of 240 placental mammals estimated by ROADIES (on the right) and the reference tree from the Zoonomia consortium ^12^ (on the left). Dashed branches show the differences between the two trees. **(C)** ROADIES convergence in accurate mode. As the number of gene trees increases, we show the percentage of highly supported species tree nodes with localPP >= 0.95 (plot i), the linear increase in runtime (ii), and the normalized Robinson Foulds distance of the final species tree to the reference tree (iii). **(D)** Quartet scores (i) and localPP branch support (ii) of all three topologies around three branches (marked in (B)), which had low support in the final tree.

#### Agreement on Atlantogenata

Historically, there has been a debate over the correct placement of the superorders Afrotheria, Xenartha, and Boreoeutheria. ROADIES aligns with Zoonomia on the hypothesis of Atlantogenata (Afrotheria + Xenartha) with full localPP support (Fig. 2B), which is also consistent with the prevailing scientific consensus ^50–56^, although alternative hypotheses of Epitheria (Afrotheria + Boreoeutheria) ^57^ and Exafroplacentalia (Xenartha + Boreoeutheria) ^58^ have also been proposed.

#### Agreement on the placement of Sirenia and Scandentia

The placement of Sirenia within Paenungulata (a clade consisting of orders Hyracoidea, Proboscidea, and Sirenia) ^57,59^ and the placement of Scandentia within the superorder Euarchontoglires have been a subject of ongoing debate ^55,57,59,60^, but ROADIES and Zoonomia tree topologies are in agreement here (Fig. 2B) and also in agreement with the growing consensus of these recent studies ^55,59,60^.

#### Disagreement within the Laurasiatheria phylogeny

ROADIES differs in the placement of Perissodactyla and Cetartiodactyla within the superorder Laurasiatheria compared to the Zoonomia reference tree, which puts Perissodactyla as the sister group to Carnivora + Pholidota (clade Zooamata). ROADIES proposes Perissodactyla as the sister group of the clade (Cetartiodactyla,(Carnivora, Pholidota)) with a modest localPP of 0.77 (topology of node 2, Fig. 2B). There is substantial gene tree discordance around this branch, with two alternatives getting quartet support above ⅓, a pattern that points to possible violations of the multi-species coalescent (MSC) model ^61^ of ILS (Fig. 2D). Since the correct placement of orders within Laurasiatheria remains contentious, with different studies suggesting Perissodactyla as the sister group to Chiroptera ^62^, Carnivora ^12,52,63^, or Cetartiodactyla ^55,64^, the inference made by ROADIES is a plausible hypothesis supported by prior studies.

#### Disagreement in the placement of Macroscelidea and Tubulidentata

There is also a difference in the placement of Macroscelidea and Tubulidentata, parts of the superorder Afrotheria, which contributes to the unresolved phylogeny of early placental mammals. Zoonomia reference tree inferred Macroscelidea as the sister of Tubulidentata ^12,29^, O’Leary et al. ^57^ inferred Tubulidentata as the sister group of Paenungulata, whereas Meredith et al. ^59^ combined Tubulidentata with Afrosoricida + Macroscelidea. ROADIES, on the other hand, recovers Macroscelidea as a sister with the clade of Tubulidentata and Afrosoricida + Chrysochloridae, albeit with a low localPP support of 0.49 (topology of node 1, Fig. 2B) and backed by only 34% of gene tree quartets (Fig. 2D). This is also a plausible hypothesis that is supported by Liu et al. ^65^, Esselstyn et al. ^55^, and Du et al. ^66^, but the low support leads us to consider this branch unresolved.

### On 240 placental mammals, ROADIES is estimated to provide over 176x speedup over conventional pipelines

The percentage of highly supported nodes increases with the gene count across the different iterations of ROADIES (Fig. 2Ci). At the end of the first iteration, the ROADIES phylogeny, built using 256 starting genes, featured 89% of branches with high support. This figure reached 97% after the fourth iteration, which utilized 4096 starting genes, marking a 0.4% change from the preceding iteration and signaling a convergence. The normalized Robinson-Foulds distances (normRF) to the reference tree kept reducing with each iteration at both species and order levels (Fig. 2Ciii), suggesting that the tree quality improved as more genes were added, though the species-level normRF appears to have leveled off in the final iteration. The total runtime of ROADIES (wall-clock time) on a single 16-core AWS instance (r6a.4xlarge) is 214h11m or roughly 9 days.

To compare ROADIES’ runtime to a conventional gold-standard pipeline, we estimated the runtime on the same instance for two of the quintessential and compute-intensive steps in the conventional pipeline: pairwise whole-genome alignments of all species using LASTZ to a reference genome (we used human genome assembly GRCh38) and the inference of orthologous regions for each of the reference protein-coding genes (we used GENCODE V38 Ensembl 104) using the state-of-the-art tool developed by the Zoonomia consortium ^12^, called TOGA ^39^, which also includes the chaining of pairwise alignments (Methods). Hence, our runtime estimate for conventional pipelines is on the conservative side since these pipelines require several other steps (such as *ab initio* gene annotation, MSA inference, gene tree estimation, and summary algorithm) that also require substantial runtime. We estimate that the two aforementioned steps (pairwise WGA and orthology inference) alone require 9513 minutes or 158h33m on a single mammalian genome-pair (*Homo sapiens* - hg38 assembly and *Mus musculus* - mm10 assembly), suggesting that the total runtime for 239 such pairs would be approximately 2273607 minutes, or 1578 days 21 hours and 27 minutes on the same 16-core AWS instance. This is not surprising, as researchers are known to spend months of wall-clock time on large compute clusters consisting of thousands of cores for phylogenetic inference at this scale ^4,34^. In other words, by eliminating compute-intensive steps of whole-genome alignments and orthology inference from the pipeline, ROADIES achieves at least 176x speedup for phylogenetic inference on this dataset while maintaining comparable accuracy to the gold-standard approaches.

#### ROADIES generalizes well on two other datasets

To evaluate the generalizability of ROADIES, we used two additional large-scale datasets consisting of (i) 100 drosophilid (pomace fly) genomes ^30^ (Supplementary Table 2), where there are relatively fewer controversies regarding the correct tree topology and (ii) the 363 avian (bird) genomes ^28^ (Supplementary Table 3), which is often regarded as one of the most challenging phylogenies within the tree of life and where the tree topology remains highly controversial. For the drosophilid dataset, we used phylogeny from the recent large-scale study by Kim et al. ^30^ as the reference. For the avian dataset, these studies in the last decade by Kuhl et al. ^67^, Prum et al. ^68^, Jarvis et al. ^4^, Feng et al. ^28^, and Stiller et al. ^6^ are considered authoritative. While these studies include all major orders of the avian phylogeny, there is significant discordance between them (Supplementary Fig. 4). We chose Stiller et al. ^6^ as our reference since it is the largest and the most recent, involving all 363 species we used in our evaluation.

#### Drosophilid dataset

On the dataset of 100 drosophilid genomes, ROADIES converged in 1105 minutes or 18h25m with 1627 gene trees (Fig. 3C). The final phylogeny had 94% high-support branches (Fig. 3Ci). Notably, ROADIES accurately identified the group-level relationships without any discrepancies with the reference (group-level normRF is 0) (Fig. 3B, C-iii), which also matches with most studies ^69–71^. ROADIES also accurately determined the relationships of species within groups, apart from a few debatable topologies within the group melanogaster as follows (species-level normRF is 0.062).

**Figure 3:**
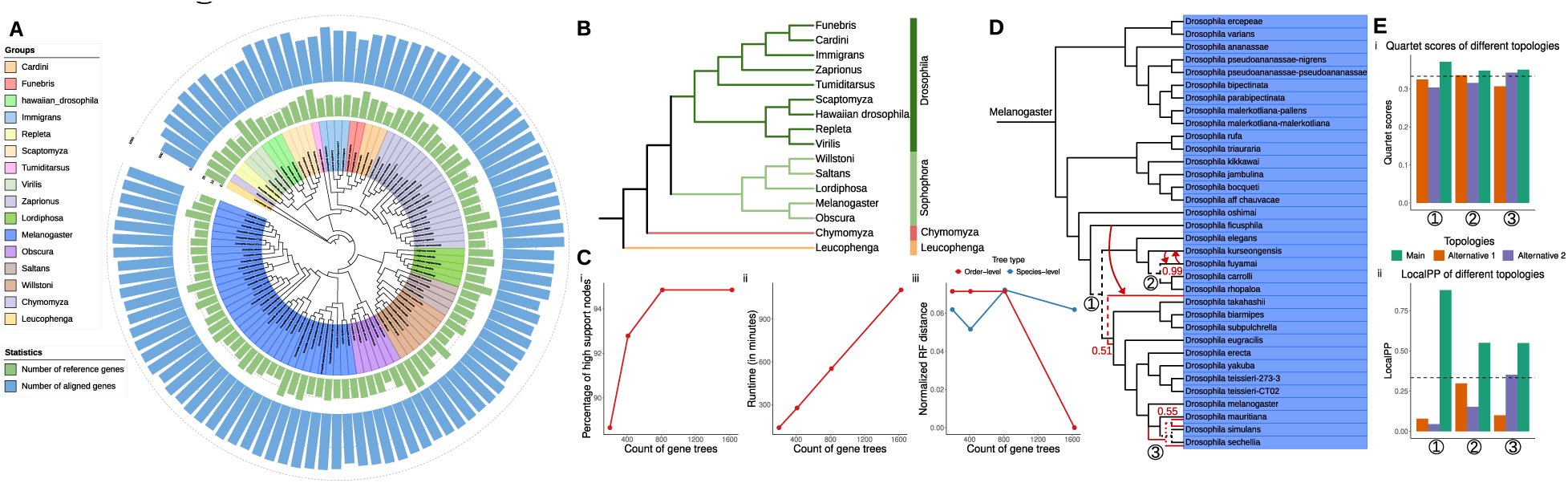
ROADIES results evaluated on the dataset of 100 drosophilid species (in the accurate mode of operation). **(A)** The species-level phylogenetic tree of 100 drosophilid species estimated by ROADIES. The number of genes aligned to each species (blue) and the count of genes sampled from each species (green) are also shown. **(B)** Group-level tree of 100 drosophilid species estimated by ROADIES, which exactly matches the reference tree ^30^. **(C)** ROADIES convergence results in accurate mode. As the number of gene trees increases, we show the percentage of highly supported species tree nodes with localPP >= 0.95 (plot i), the linear increase in runtime (ii), and the normalized Robinson Foulds distance of the final species tree to the reference tree (iii). **(D)** Species-level tree from the group melanogaster, estimated by ROADIES. Low-confident branches that differ from the reference tree are highlighted. The ones highlighted in red are from the reference tree. **(E)** Quartet scores (i) and localPP branch support (ii) of all three topologies around three branches (marked in (D)), which had low support in the final tree.

##### Differences in the positions of Drosophila mauritiana, Drosophila simulans and Drosophila sechellia

Kim et al. ^30^ infers *Drosophila mauritiana* and *Drosophila simulans* as sisters with the relatively lower support value of 0.55 (local posterior probability used by Kim at al. ^30^ is reported by ASTRAL) (topology of node 3 highlighted in red, Fig. 3D), compared to the rest of the branches. ROADIES, on the other hand, infers *Drosophila sechellia* and *Drosophila simulans* as sisters with a support value of 0.548 (main topology of node 3, Fig. 3Eii). The low support values for the main topology suggest high discordance in the gene trees, as also evident from the relatively close Quartet scores for the different alternatives (Fig. 3Ei).

##### Differences in the position of Drosophila kurseongensis

Kim et al. ^30^ places *Drosophila kurseongensis* as a sister branch with the clade of (*Drosophila carrolli*,*Drosophila rhopaloa*) with a support value of 0.99 (topology of node 2 highlighted in red, Fig. 3D). On the other hand, ROADIES infers *Drosophila kurseongensis* as a sister with the clade of (*Drosophila fuyamai*, (*Drosophila carrolli*,*Drosophila rhopaloa*)) with localPP of 0.549 (main topology of node 2, Fig. 3Eii). The difference in the node 2 topology inferred by ROADIES compared to the reference tree might be related to sampling or statistical inconsistency since similar ROADIES experiments (refer to the section “ROADIES is scalable and stable”) inferred the exact topology with the reference tree with a localPP of 1.

##### Differences in the position of Drosophila ficusphila

The placement of *Drosophila ficusphila* by ROADIES is supported by localPP of 0.876 (main topology of node 1, Fig. 3Eii), whereas the reference tree infers a different position with the confidence score of 0.51 (topology of node 1 highlighted in red, Fig. 3D). These differences indicate that topologies with lower confidence in the reference tree are more likely to be contested and are also represented differently by ROADIES, suggesting the need for further research.

#### Avian dataset

ROADIES required 2811 instance hours on a single 16-core AWS EC2 instance (r6a.4xlarge) to reach convergence for 363 avian genomes dataset (we parallelized the pipeline using 16 such instances to ensure it completed within a reasonable timeframe). The final species tree consisted of 99% highly supported nodes. Since the rapid radiation of birds presents a challenging case, it took much longer – 7 iterations with a total of 62,413 gene trees – for the topology to stabilize and reach our convergence criteria (Fig. 4Ci). Remarkably, this number is also close to the 63,430 gene trees Stiller et al. ^6^ used in their analyses. The converged ROADIES phylogeny achieved a normRF score of 0.027 compared to our reference tree at the species level (Fig. 4Ciii), suggesting a reasonably high congruence between the two phylogenies. Here, too, every species was correctly assigned to its respective phylogenetic order, and the differences only existed at the order level and in the organization of species within those orders, mostly among low support branches (Fig. 4A-B).

**Figure 4:**
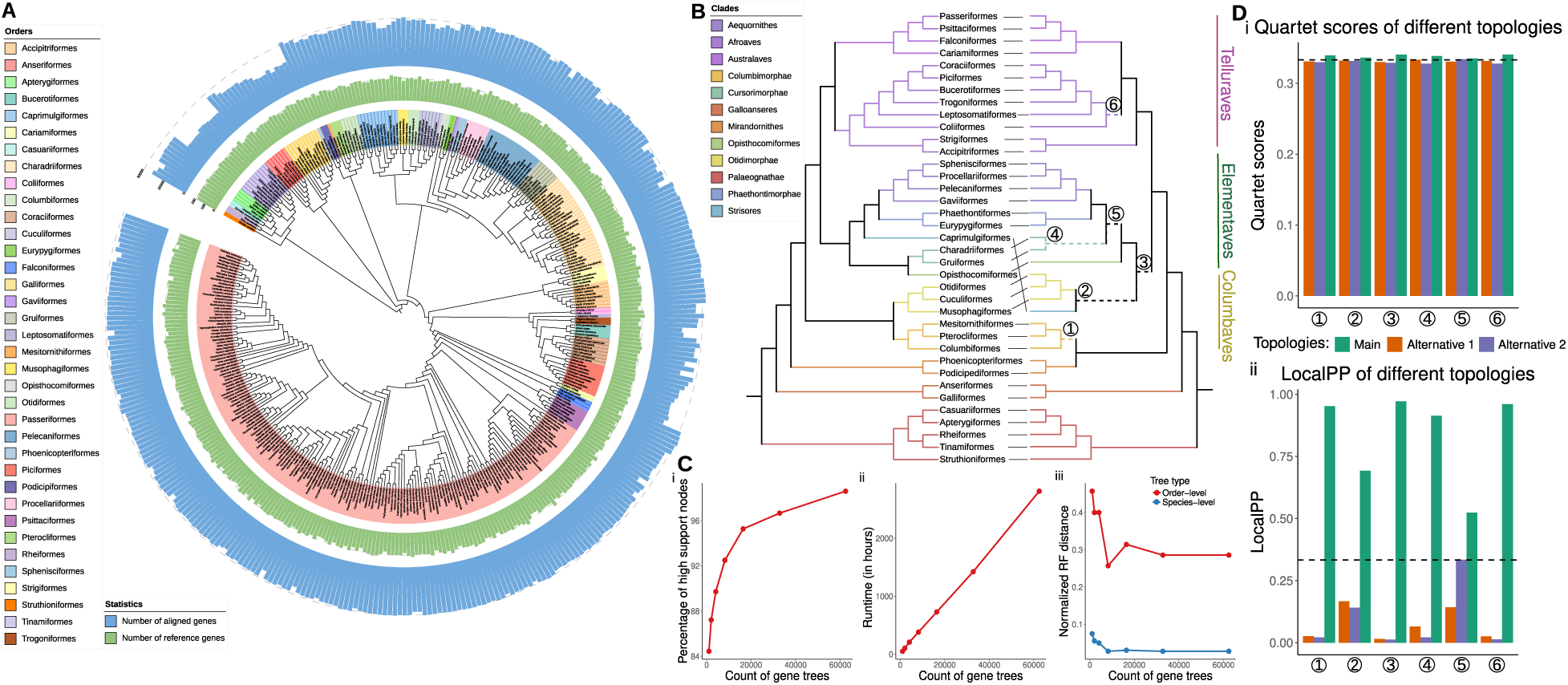
ROADIES results evaluated on the dataset of 363 avian species (in the accurate mode of operation). **(A)** The species-level phylogenetic tree of 363 avian species estimated by ROADIES. The number of genes aligned to each species (blue) and the count of genes sampled from each species (green) are also shown. **(B)** Order-level trees of 363 avian species estimated by ROADIES (on the right) and the reference tree from Stiller et al. ^6^ (on the left). Dashed branches show the differences between the two trees. **(C)** ROADIES convergence results in accurate mode. As the number of gene trees increases, we show the percentage of highly supported species tree nodes with localPP >= 0.95 (plot i), the linear increase in runtime (ii), and the normalized Robinson Foulds distance of the final species tree to the reference tree (iii). **(D)** Quartet scores (i) and localPP branch support (ii) of all three topologies around six branches (marked in (B)), which had low support in the final tree.

At the order level, ROADIES-inferred phylogeny captured known relationships but also had differences with the reference tree among hard to resolve branches. Overall, six ordinal-level branches were different, and all of those are among those that are contentious, and in every case, these had quartet frequencies that were close to ⅓, which signifies a polytomy (Fig. 4D). ROADIES and the reference tree are aligned in (i) capturing the relationship of orders within Australaves and Aequornithes within Neoaves and (ii) in the relationship of orders within core waterbirds, both of which have been involved in considerable debate, and have been even suggested by some autors to represent a hard polytomy ^72^.

##### Differences in Palaeognathae

The placements of Tinamiformes and Rheiformes inferred by ROADIES differ from those in the reference tree, but past studies have not reached a consensus in this case. Some studies support ROADIES’ findings ^73,74^, whereas some propose other topologies^75^. For this relationship, Stiller et al. ^6^ presented a CoalHMM analysis supporting the same resolution found by ROADIES, which led them to treat the resolution of this node as uncertain.

##### Early Neoavian diversification

ROADIES infers Columbimorphae and Phoenicopterimorphae as sister groups, recovering the Columbea group proposed by Jarvis et al. ^4^ and Yuri et al. ^75^ but not in our reference tree ^6^. As Mirarab et al. ^76^ have shown, this result is likely due to a region of chromosome 4 of chicken, with an unusually strong signal for Columbea across 21Mb. The fact that this region has a unique history across such long stretches conflicts with the MSC model and was conjectured to be due to complex rearrangement polymorphisms persisting through multiple speciation events. Mirarab et al. ^76^ also showed that the presence of this recombination-depleted region makes the inferred species tree sensitive to the choice of loci. Related to this change is another one: ROADIES infers Otidimorphae as sisters to Caprimulgiformes, breaking the Elementaves group proposed by Stiller et al. ^6^. However, we note that the support for this relationship in ROADIES is low (localPP=0.69) (Fig. 4D), and the ROADIES tree is compatible with Elementaves if we contract branches of localPP < 0.95.

##### Other differences

ROADIES also disagreed with the reference trees in two relationships where Stiller et al. ^6^ had relatively low confidence in their results due to sensitivity to data subsampling: ROADIES puts Accipitrimorphae as sister to all other land birds (Telluraves), a position that Stiller et al. did not fully rule out. Lastly, Opisthocomiformes, the only order with low support in the Stiller et al. analyses, also moved within Elementaves. Neither tree is confident in its placement of this rogue taxon.

Overall, the much slower convergence of ROADIES matches the understanditng that avian phylogeny is among the most challenging. Nevertheless, the tree found is similar to the set of trees that have been identified in recent genome-wide analyses, recovering the so-called magnificent seven groups ^77^. Finally, the support values are low for the branches that differ between our tree and the reference tree.

### Comparison to another fully automated approach: MashTree

There have been decades of research in the area of phylogenetics, yet we are aware of only one computational tool, MashTree ^24^, that is capable of directly inferring species trees from raw genome assemblies, underscoring the challenging nature of this problem. Relative to ROADIES, MashTree is a relatively simplistic pipeline involving just two main steps: (i) calculating the evolutionary distances between different genome assemblies with Mash ^78^ and (ii) leveraging the Mash distance matrix to deduce the species tree through the Neighbor-Joining algorithm, as implemented in QuickTree ^79^. To compare the performance and accuracy of MashTree with ROADIES, we evaluated it on all three datasets (mammals, pomace flies, and birds) (Fig. 5A-C). Unsurprisingly, we found that MashTree was much faster than ROADIES on all datasets. However, it also had significantly higher normRF distances to the reference phylogeny, signaling accuracy issues (Fig. 5D). For example, in the drosophilid phylogeny, which is relatively stable and has a broad scientific consensus, MashTree’s tree was inferred in only 1 minute but had a group-level distance of 0.14 relative to the reference. Notably, MashTree inaccurately inferred orders Obscura and Melanogaster within the Drosophila subgenera. Additionally, it inferred an implausible placement for the order Tumiditarsus (Fig. 5A).

MashTree’s order-level distances for the mammalian and avian phylogenies were also significantly worse, 0.61 and 0.714, respectively, with startling inaccuracies even within well-established clades, such as the positioning of superorders Xenartha, Afrotheria, and Boreoeutheria in mammals. MashTree also struggled to distinguish between Euarchontoglires and Laurasiatheria within Boreoeutheria, erroneously placing Eulipotyphla with Euarchontoglires. Additionally, the arrangement of orders within Laurasiatheria and Afrotheria differs between MashTree and the reference tree, while ROADIES aligns more closely with the reference (Fig. 5B). For birds, MashTree-inferred order-level tree topology seems almost random, with little alignment to the reference tree across the majority of orders (Fig. 5C).

In summary, although MashTree may be effective at categorizing species into broad phylogenetic orders, it does not appear to have the accuracy in determining the relationships among those orders or the relationships of species contained within those orders, which is necessary for broad scientific applicability. For these reasons, we believe ROADIES could be the first fully automated approach to species tree inference that has broad biological and medical utility due to its high overall accuracy.

### ROADIES offers flexibility in balancing speed and accuracy

Species tree inference often involves a trade-off between the speed and accuracy of the final results. In most biological applications, accuracy is paramount, though speed is desirable and often a constraint. We have implemented ROADIES as a Snakemake workflow ^80^, which allows modular implementation and exposes numerous parameters of individual tools to end users (Methods). This is to allow expert users the flexibility to fine-tune the parameters of individual stages to optimal settings depending on the nature of the workload. It is also easy to swap tools in particular stages of the ROADIES pipeline with equivalents, such as substituting RAxML-NG ^44^ with IQTree^81^. We anticipate that most users will not adjust the ROADIES pipeline to such an extent and will likely prefer the default mode prioritizing accuracy. However, we have provided two convenient additional modes in ROADIES, fast and balanced, to cater to users who might be resource-constrained or might be working with datasets or applications that might be more tolerant of errors (see Methods for details).

We analyzed the 240 placental mammals dataset with balanced and fast modes of operation in ROADIES to evaluate them in terms of accuracy and efficiency. Supplementary Fig. 1 highlights these results. In balanced mode, ROADIES required 3738 gene trees to reach convergence, with 96% of high support nodes in the final tree and consuming 160h55m on a 16-core AWS EC2 instance (r6a.4xlarge) (Supplementary Fig. 1Ei). In fast mode, the convergence was slower – requiring 14929 gene trees – but the overall runtime was still lower, 98h21m on the same instance. The fast mode resulted in 97% of high support branches in the final species tree (Supplementary Fig. 1Fi). Overall, the balanced and fast modes on this dataset were 1.33x and 2.17x times faster than the accurate mode, respectively. In terms of accuracy, both balanced and fast modes provided a more accurate tree than MashTree ^24^, with substantially lower species-level and order-level normRF distances from the reference tree (Supplementary Fig. 1Eiii,1Fiii). Notably, both fast and balanced modes accurately estimated the once-highly debated relationship of three superorders Afrotheria, Xenartha, and Boreoeutheria (with a clear split between Euarchontoglires and Laurasiatheria within Boroeeutheria) ^50–56^, (Supplementary Fig. 1A-B). The balanced mode also captured the same relationship of orders within Afrotheria as the reference, resulting in a slightly better order-level distance than the accurate mode. There is only one difference in the arrangement of orders in the balanced mode compared to the reference tree – the placement of Perissodactyla and Cetartiodactyla (Supplementary Fig. 1C). In the fast mode, there are a few extra differences on top of the changes for balanced mode, such as the placements of Afrosoricida and Tubulidentata ^55,57,59,65^, the placement of Scandentia in Euarchontoglires ^55,57,59,60^, the placement of Perissodactyla and Cetartiodactyla in Laurasiatheria ^52,55,57,62–64^, and the placement of Proboscidea in the clade Paenungulata ^57,59^ – each of these differences has been the subject of previous debates (Supplementary Fig. 1D).

We also analyzed the phylogeny of pomace flies and birds with these two modes of operation (Supplementary Fig. 2 and 3, respectively). In the balanced mode for pomace flies, ROADIES required 8192 gene trees to reach convergence with 100% highly-supported nodes in the final tree and a runtime of 16h17m on a 16-core AWS EC2 instance (r6a.4xlarge) (Supplementary Fig. 2Ei). In the fast mode, convergence was reached with 4096 gene trees, with a total runtime of 1h59m on the same instance and 95% highly-supported branches in the final species tree (Supplementary Fig. 2Fi). Both balanced and fast modes successfully captured highly stable orders like Chymomyza, Leucophenga, and certain stable orders within the subgenera Drosophila and Sophophora ^69–71^. In the balanced mode, only the groups Melanogaster and Obscura were incorrectly placed within the subgenera Drosophila, whereas the reference tree placed them within the subgenera Sophophora (Supplementary Fig. 2C). In the fast mode, in addition to the aforementioned change, there were differences in the placement of orders Tumiditarsus and Zaprionus within the subgenera Drosophila (Supplementary Fig. 2D). For birds, in the balanced and the fast mode, ROADIES required 62441 gene trees with 98% and 31692 gene trees with 96% respectively, with runtimes of 1066 hours and 101 hours (Supplementary Fig. 3). Overall, on this dataset, the balanced and fast modes were 2.59x and 27.83x times faster than the accurate mode, respectively, but with lower accuracy.

To summarize, the accuracy of both balanced and fast modes seems to be reasonably high and may be sufficient for some applications. We note that the fast mode in ROADIES combines a summary method with distance-based gene tree estimation, and our results showed this approach has higher accuracy than a genome-wide distance-based species tree estimation (MashTree, Supplementary Fig. 5).

### ROADIES is scalable and stable

We have designed ROADIES to support large-scale phylogenetic analyses that several future applications would necessitate. Such analyses are typically performed on high-performance computing (HPC) environments, requiring days to months of processing time after distributing the workload across several compute servers. Since ROADIES has been implemented using Snakemake ^80^, it allows convenient and efficient parallelization of the workflow in different computing environments, including the cloud platforms. Specifically, ROADIES is able to easily exploit parallelism at the thread and process levels. To characterize the strong scaling efficiency of ROADIES, we ran ROADIES with varying numbers of system cores in the AWS EC2 instance for 100 drosophilid datasets in accurate mode (Methods). We fixed the gene count to 4000 for these experiments so that variance in the convergence rate does not affect the runtime estimates. We observed roughly linear speedup with the number of cores – the runtime decreased from 3147 minutes to 430 minutes as the number of cores was increased from 8 to 128, achieving a scaling efficiency of 57.7% (Supplementary Fig. 6A). The slight increase in the runtime with 256 cores is primarily due to the communication overhead, which we will address in the future releases. Additionally, we investigated how the runtime scaled as we varied the number of input species, keeping the core count fixed at 16, and the gene count at 4000. The runtime increased from 90 to 1778 minutes as the number of species went from 15 to 100, indicating a super-linear but sub-quadratic scaling in runtime with respect to the number of species (Supplementary Fig. 6B).

Given that ROADIES uses random sampling of genes from input genomes, it could be susceptible to sampling noise. To assess the consistency of the phylogenetic trees produced by ROADIES, we conducted four separate trials on the drosophilid dataset, allowing ROADIES to randomly select a separate set of genes at each run. We observed that one experiment (experiment ID 3, Supplementary Fig. 6E) converged faster in 4 iterations compared to 5 iterations in the rest. The final phylogenies produced by all experiments were nearly identical, with the maximum normRF distance between any two trials being only 0.04 at the species-level (Supplementary Fig. 6C-F). Moreover, the percentage of highly supported nodes in the final converged species tree did not differ much across all four experiments and followed a similar trend with increasing gene tree count (Supplementary Fig. 6E).

The slight differences in species-level normRF distances between pairs of unique experiments result from a few debatable drosophilid species phylogenies: i) *Drosophila fuyamayi* and *Drosophila kurseongensis* - where some experiments align with our reference, whereas some differs with low support (localPP = 0.587 and 0.652), ii) *Drosophila mauritiana*, *Drosophila simulans,* and *Drosophila sechellia* - where all experiments differ with the reference. Even reference tree proposes this topology with lower support values (localPP = 0.55), and iii) *Drosophila quadrilineata* and *Drosophila repletoides* – where 3 out of 4 experiments match the reference. This shows that ROADIES experiments are mostly in consensus with each other, barring a few debatable and low-confident topologies as exceptions. In addition, the order level normRF distance is consistent with each other, except one group Tumiditarsus (representing *Drosophila repletoides)*, which makes order level normRF jumps from 0 to 0.07 in a few cases. Given these observations, we conclude that the ROADIES phylogenies are stable, with negligible dependence on the specific genes it samples.

## Discussion

ROADIES is the first-of-its-kind tool that can perform discordance-aware species tree construction directly from raw genome assemblies without relying on a single reference genome, input alignments, gene annotations, and orthology inference. With genomic data continuing to grow exponentially, and with telomere-to-telomere assemblies now possible for complex genomes at reasonable costs ^12,28,82^, ROADIES is addressing an urgent and critical problem and is likely to find diverse applications in biological and medical fields. For large datasets, ROADIES can provide up to two orders of magnitude improvement in computational costs compared to conventional methods while achieving comparable accuracy. ROADIES achieves these impressive results based on three key insights. *First*, instead of genomic regions with specific characteristics, such as protein-coding genes, ultra-conserved elements (UCEs) ^83^, or transposons ^63,84–86^. ROADIES is based on a random sampling of loci from input genomes. This eliminates the need for genome annotations and the bias caused by using a single reference genome. *Second*, ROADIES uses recent developments in summary methods for species tree inference that can work with multicopy gene trees that account for both orthology and paralogy ^37^. This eliminates the need for strict orthology inference – a longstanding problem in computational genomics. While good progress has been made on this problem in recent years, such as the development of TOGA ^39^, orthology inference typically requires whole-genome alignments and other intensive algorithms to accurately distinguish orthologs from paralogs and pseudogenes. On the other hand, homology (combination of orthology and paralogy) is not just easier to infer; it is also computationally inexpensive and less error-prone. *Third*, choosing the correct number of gene trees is an essential challenge in summary methods, as it heavily depends on the complexity of the evolutionary histories of the input genomes, which is often unknown in advance. ROADIES tackles this issue through a convergence algorithm that initiates with a small set of genes, doubles the number in each subsequent iteration, and stops once the difference in topological support (measured as the percentage of branches with high support) between two successive iterations becomes minimal. As expected, ROADIES requires a far greater number of gene trees to converge for bird phylogeny, the most complicated case in our datasets, compared to the phylogeny of mammals and flies.

In the future, we aim to further improve the scalability, accuracy, and capabilities of ROADIES. While our current implementation is likely to work well with thousands of vertebrate or mammalian genomes, the runtime might be unreasonably high for tens of thousands of genomes or beyond. We believe GPUs could be leveraged to accelerate the critical stages of our pipeline ^87,88^, but this would necessitate the development of new GPU-accelerated libraries. Moreover, incorporating recently proposed divide-and-conquer strategies for species tree construction also presents a promising avenue for improving ROADIES ^89^. Our analyses encompassing mammals, flies, and birds, have been confined to evolutionary timescales of approximately 5–100 million years ^6,12,30^. Expanding our approach to significantly longer or shorter timescales poses substantial challenges. For longer timescales, the issue of sequence divergence reaching the ‘twilight zone’ of sequence alignment becomes a significant hurdle ^90^, making it difficult to establish reliable homologies. Conversely, for shorter timescales, the strategy of random gene sampling may not consistently capture phylogenetically informative sites ^91^. Lastly, we are exploring advanced algorithms for tree rooting ^92,93^ and methods for quantifying phylogenetic uncertainty ^94,95^.

## Methods

### Input Datasets and Reference Phylogenies

Three genomic assembly datasets for recent large-scale studies were used for evaluation in this manuscript consisting of (1) 240 species from the infraclass Placentalia (alternatively referred to as “placental mammals”), (2) 100 species belonging to the subfamily of Drosophilinae and Steganinae, and (3) 363 species from the class Aves. The details of these genomic datasets, including their respective orders, scientific nomenclature, and NCBI ID, are provided in Supplementary Table 1-3, respectively. The 240 species of placental mammals are collected from the Zoonomia consortium ^12^. Zoonomia consortium earlier published 241 species with two assemblies from the same species - *Canis lupus familiaris* (Domestic dog). We removed the duplicate dog assembly (with NCBI ID GCF_000002285.3) and kept the one with NCBI ID GCA_004027395.1. The 363 avian species are collected from the Birds 10k Genome Project’s dataset ^28^. For 100 drosophilid genomes, we collected the dataset from NCBI BioProject ID PRJNA675888 used by Kim et al. ^30^.

We used reference trees from authoritative studies as a proxy for ground truth to compare the accuracy of the species tree estimated by ROADIES. The reference tree of 240 placental mammals is taken from Zoonomia consortium ^12^. The reference tree of 100 drosophilid genomes is taken from Kim et al. ^30^. The reference tree of 363 Avian species is collected from the topology described by Stiller et al. ^6^.

### ROADIES: Implementation Details

ROADIES involves multiple stages of processing (Fig. 1) and has been implemented as a Snakemake workflow for modularity. It starts with a random sampling of subsequences from input genomic sequences where each subsequence is considered a “gene”. Next, it performs pairwise alignment through LASTZ ^96^ to find homologs of all sampled genes within the input genomic sequences. Then, it filters out low-quality alignments to prevent redundant computation in subsequent stages and performs multiple sequence alignments of homologous genes using PASTA^43^. Next, it estimates gene trees from multiple sequence alignments of genes using RAxML-NG ^44^, and at the last stage, constructs a species tree using ASTRAL-Pro2 ^37^ from those gene trees. The implementation details of each stage are described in individual subsections below. Every stage in ROADIES can be customized easily by altering the default parameters in a single configuration file named `config.yaml`.

#### 1) Random Sampling of Genes

For executing the random sampling approach, ROADIES takes raw genomic sequences of multiple species, each in `.fa` format or `.fa.gz` format as input and randomly samples a specified number of subsequences or genes (the number of samples is parameterized as GENE_COUNT in the configuration file) of a fixed size (the size of each sampled gene is parameterized as LENGTH) across all input genomes. For our analysis in this manuscript, we configured LENGTH as 500 (each gene of length 500 base pairs; we experimented with varying gene lengths from 250 to 1000 and observed 500 to provide the best results in terms of accuracy and speed).

In this random sampling approach, each sampled subsequence is treated as a gene. ROADIES iterate through all input genomes to get the desired number of genes. All sampled gene sequences are assigned a unique ID number to be represented in later stages and are saved in a common `out.fa` file. Along with random sampling, ROADIES filters and discards the low-quality genes by setting a threshold on the percentage of accepted aligned (i.e., upper-case) characters in a gene, configured by the UPPER_CASE parameter (default UPPER_CASE is 0.9 to discard the genes that have less than 90% of accepted upper-case characters). Genomic sequences with ambiguous bases (i.e., not “A”, “C”, “G”, and “T”) and lower-case characters are considered noisy, and we discard them.. ROADIES also plots the number of genes sampled from each genome across the entire input dataset for easy analysis (Supplementary Fig. 7A).

#### 2) Pairwise Alignment to find Homologs

In the second stage, ROADIES takes a list of sampled genes from the previous random sampling stage and performs the pairwise sequence alignment of all genes with all genomic reference assemblies individually through the state-of-the-art tool LASTZ (version 1.04.15) ^96^. The default LASTZ command that is incorporated in ROADIES is:

`lastz_32 {input genomes path}[multiple] {input genes path} -- coverage=85 --continuity=85 --filter=identity:65 --format=maf --step=1`

`--queryhspbest=100.`

LASTZ takes one unique genomic sequence and a list of sampled gene sequences from the out.fa as input and performs pairwise alignment of each genome with all sampled genes to find the homologous sequences.

Among LASTZ parameters, `--continuity` defines the allowable percentage of the non-gappy alignment columns and discards the alignments with more gaps. The high value of continuity is preferable to remove long, gappy fragmented alignments since gappy alignments slow down the computation of multiple sequence alignments, less likely to be true homologs, and fragments can reduce accuracy of gene trees ^97^. ROADIES has the default value of `--continuity` set as 85, which is user-configurable through the CONTINUITY parameter. `--identity` defines the percentage of aligned matched bases. ROADIES sets the default `--identity` value as 65, user-configurable through the IDENTITY parameter. Lastly, ROADIES has the default value of `-- coverage` as 85, which is user-configurable through the COVERAGE parameter, which sets the threshold for the percentage of the input sequence to be included in the alignment block.

LASTZ performs pairwise alignment, which is repeated for all input genomic sequences to get the homologous sequences across all genomes in the input dataset. The output alignment from LASTZ for each input genomic sequence are saved as a `<SPECIES_NAME>.maf file`. `<SPECIES_NAME>.maf` file contains information about the alignment score, positions, and gap penalties for each pairwise alignment with a gene for all genomes, helping ROADIES filter low-quality alignment at the next stage. Supplementary Fig. 7B shows the number of genes (sampled subsequences) aligned to each genomic data in the entire dataset after the second stage of ROADIES.

#### 4) Filtering Low-Quality Alignments

Some pairwise alignments from the previous step can be gappy or uninformative to construct accurate phylogenetic trees. These extra gappy alignments increase computational burden without adding much to the accuracy; hence, it is crucial to filter them out before proceeding to further stages. To execute this filtration of low-quality alignments, ROADIES collects the pairwise alignments from each species’ `<SPECIES_NAME>.maf` file, collects the gene-specific alignment information, and stores the alignments belonging to a particular gene in a separate `gene_<id>.fa` file where the `id` corresponds to the ID number of that particular gene. This process is done for all genes across all `<SPECIES_NAME>.maf` files.

Once alignments are gene-wise sorted, ROADIES performs multiple levels of filtering to remove low-quality gappy alignments from each `.maf` file. Firstly, ROADIES sets a threshold to allow only up to a maximum number of high-scoring alignments from each type of gene and from the same species, configured by the MAX_DUP parameter (default MAX_DUP is set to 10 to allow only ten high-scoring alignments from a type of gene from the same species). `gene_dup.png` in the *plots* folder of the specified OUT_DIR shows the histogram plot of the duplicate gene count (Supplementary Fig. 7C). ROADIES also prevents the occurrence of several alignments sampled from a very small genomic region by discarding the alignments whose starting positions are less than 2000 base pairs. For example, four alignments with starting positions within a genomic region (10,000-10,500) at positions 10,000, 10,070, 10,309, and 10,249 are prevented and filtered out after keeping only one alignment with the maximum score. Secondly, ROADIES sets a minimum threshold of the number of allowed species for a particular gene to proceed with the following stages, configurable through the MIN_ALIGN parameter. If a gene is rare and found in fewer species than the threshold (i.e., four species - default MIN_ALIGN value), it will not help build accurate phylogenetic trees. Thus, those genes are filtered out.

#### 5) Multiple Sequence Alignment of Filtered Genes

After filtering uninformative alignments in the previous stage, ROADIES proceed to its fourth stage, which performs multiple sequence alignment (MSA) across all alignments stored in a particular gene fasta file (`gene_<id>.fa`). This process is executed for all genes. ROADIES performs MSA using the state-of-the-art tool PASTA (version 1.9.0) ^43^ (`python pasta/run_pasta.py -i {input gene fasta file path} -j “gene_{id}” --alignment-suffix=”.fa.aln” --num-cpus {number of threads}`). All the MSA outputs are saved as `gene_id.fa.aln` file. Additionally, before executing MSA for a gene fasta file, ROADIES checks if the number of filtered alignments in that gene fasta file is more than or equal to MIN_ALIGN (default value of MIN_ALIGN is 4).

After MSA computation, ROADIES perform further filtration of gappy MSAs and mask out gappy sites from MSAs through a script `run_seqtools.py` included with PASTA (`python run_seqtools.py -filterfragmentsp 0.5 -masksitesp 0.02 -infile {input MSA file path} -outfile {output filtered MSA file path}`). PASTA provides multiple options to filter out the gappy alignments across the columns and rows with `run_seqtools.py. --filterfragmentsp` removes the gappy alignment rows from MSA with less than the specified portion of the non-gap characters. `--masksitesp` masks out gappy sites across columns of the MSA having less than the specified portion of the non-gap characters. ROADIES sets mask gappy sites threshold (`--masksitesp`) to 0.02, allowing sites to have up to 98% of the gaps; this minimal filtering manages to remove many gappy sites, which can result from PASTA alignments.It sets filter fragment threshold (`--filterfragmentsp`) as 0.5 (filtering out alignments with 50% gaps) following suggestions from previous studies ^97^. Users can configure these options through FILTERFRAGMENTSP and MASKSITESP parameters in `config.yaml` file.

In a few cases, all sequences in a gene fasta file can be identical, which will make them unsuitable for gene tree estimaiton. ROADIES prevent such a scenario by keeping a check on identical alignments and, if they exist, bypassing the execution of this stage and proceeding directly with the next stage of gene tree estimation, which will eliminate these cases.

#### 6) Gene Tree Estimation

In its fifth stage of execution, ROADIES collects all the multiple sequence alignment outputs from the previous stage and performs gene tree estimation using RAxML-NG ^44^ (`raxml-ng --msa {input msa file path} --model GTR+G+F --redo --threads auto{{{number of threads}}} --blopt nr_safe`). Before executing the gene tree estimation step, ROADIES checks if the number of unique species in an input filtered MSA file (which belongs to a particular gene) is more than or equal to four (default value of MIN_ALIGN parameter). If it is less than four, further computation will not be executed.

The output file with the estimated gene tree will be saved as `gene_<id>.filtered.fa.aln.raxml.bestTree`. This process is repeated for all the genes, and in the end, it concatenates all treefiles from individual genes into a common file (`gene_tree_merged.nwk`) containing the list of gene trees, which is then given to ASTRAL- Pro2 ^37^ at the next stage for species tree estimation. The total number of gene trees in the concatenated list will be closer to the actual configured GENE_COUNT parameter (but may not be the same because of the filtration strategy in the intermediate stages, which filters out some of the genes - Supplementary Fig. 7D) For gene IDs containing identical homologous sequences, RAxML-NG would skip the tree estimation. Hence, there will not be any inferred gene trees for that gene ID.

Among the DNA substitution models implemented in RAxML-NG ^44^, we chose the General Time Reversible model (GTR) with a discrete Gamma model having default four rate categories (+G) and with empirical base frequencies (+F).

#### 7) Species Tree Estimation

This is the last stage of our pipeline, where ROADIES takes the list of gene trees from the previous stage as input (`gene_tree_merged.nwk`) and performs the species tree estimation step using ASTRAL-Pro2 ^37^ (`ASTER-Linux/bin/astral-pro2 -i {input gene tree list file path} -o {output species tree} -a {mapping file path}`). Along with the list of gene trees, ASTRAL-Pro2 requires a mapping file containing the list of names of the genes mapping to their corresponding species name. This file is generated at the filtering stage (`mapping.txt`) and is fed as input to this stage. ASTRAL-Pro2 creates a single species tree in the Newick format.

ASTRAL-Pro2 also estimates local posterior probability (localPP) support for branches of the final species tree by examining quartet frequencies around each branch, which is also outputted () in a file called `freqQuad.csv` (`ASTER-Linux/bin/astral-pro2 -u 3 -i {input gene tree list file path} -o {output species tree} -a {mapping file path}`). ROADIES thus keep track of the percentage of highly supported nodes (>0.95) in the final species tree, estimating the confidence of the tree (in `--converge` mode - refer to section 4.11). In addition to confidence values, ASTRAL-Pro2 also estimates the branch lengths in the final Newick tree saved as `roadies.nwk` and provides the localPP of all three topologies in each node of the final species tree, saved as `roadies_stats.nwk`. ASTRAL-Pro2 provides unrooted species trees by default. Hence, users are required to root the output tree.

##### Modes of Operation

ROADIES supports various modes of operation, including accurate, balanced, and fast mode, which offers flexible runtime-accuracy tradeoffs to the user. Out of all the stages in the pipeline, gene tree estimation is the most compute-intensive and time-consuming stage (Supplementary Fig. 8). Hence, we targeted to reduce the runtime of the overall pipeline by optimizing this stage.

##### Accurate mode

Accurate mode is the default mode of operation where multiple sequence alignment and gene tree estimation stages of ROADIES are governed by PASTA ^43^ and RAxML-NG ^44^, respectively, as mentioned in the above sections.

##### Balanced mode

In the balanced mode of operation, multiple sequence alignment is performed by PASTA ^43^, and the gene tree estimation is performed by FastTree (version 2.1.11) ^45^. The execution of multiple sequence alignment is similar to the accurate mode of operation. For gene tree estimation, the msa output from the PASTA in the form of `gene_{id}_filtered.fa.aln` is provided to FastTree ^45^ (`FastTree -gtr -nt {input msa file path} > {output file path}`) for estimating gene trees from the given msa. The gene tree output of the FastTree (`gene_{id}_filtered.fa.aln.treefile`) is concatenated for all gene IDs and provided to ASTRAL-Pro2 ^37^ for final species tree estimation. Before estimating gene trees by FastTree, ROADIES checks the presence of MIN_ALIGN alignments and discards the execution of the gene ID having less than MIN_ALIGN alignments. FastTree is considered to be relatively faster than IQTREE-2 ^81^ at the expense of accuracy ^98^. In this mode, users expect a shorter runtime than in accurate mode, and it can be used in runtime-critical scenarios.

##### Fast mode

In the fast mode of operation, the multiple sequence alignment and the gene tree estimation stages are performed together by Mashtree ^24^, which uses distance-based tree estimation methods. In this mode, ROADIES takes individual gene fasta files (`gene_<id>.fa`) and checks if the number of filtered alignments in that gene fasta file is more than or equal to MIN_ALIGN (default value of MIN_ALIGN is 4). If it has more than MIN_ALIGN alignments, ROADIES estimates the gene trees by executing MashTree (`mashtree --mindepth 0 --numcpus {number of threads} --kmerlength 10 {input_gene_file_path}/*.fa > {output_path}`). The output of the MashTree in the form of `gene_{id}.dnd` for all gene IDs is concatenated and fed to ASTRAL-Pro2 ^37^ for species tree estimation.

##### ROADIES Convergence Mechanism

ROADIES follows a convergence mechanism for estimating a stable and confident species tree. Here, ROADIES iterates multiple times, where, at each subsequent iteration, the number of genes gets doubled. The list of gene trees from all previous and current iterations together gets concatenated at every iteration, which is provided to ASTRAL-Pro2 ^37^ for final species tree estimation. At every iteration, ROADIES evaluates the percentage of high support/confident nodes (nodes with localPP >= 0.95) in the final species tree (`roadies_stats.nwk`). ROADIES continues iterating until it reaches the convergent point where the change in the percentage of high support nodes in the current iteration from the previous iteration is less than 1% and stops further execution. The final species tree at the last iteration is considered to be the converged species tree (Figure 1C). In the entire convergence process, the list of gene trees obtained from previous iterations gets iteratively concatenated, resulting in an expanding set of gene trees. In the convergence mode, the starting count of the genes can be parameterized as GENE_COUNT, from which it gets doubled at every iteration. The more the genes are, the more stable and accurate the final trees will be.

##### ROADIES Snakemake Workflow

ROADIES pipeline execution uses Snakemake workflow ^80^ within a custom Conda ^99^ environment where the prerequisite tools needed for each stage in the pipeline are installed and added. All the stages of ROADIES are independent snakemake rules, with corresponding input and output files and the required parameters. The snakemake workflow consisting of the set of rules as individual stages of the ROADIES pipeline is called using the command: `snakemake --cores (number of cores) --config mode=(accurate/balanced/fast) config_path=(input config path) num_threads=(number of threads) --use-conda --rerun- incomplete`.

ROADIES provides the benefit of both multithreading and multiprocessing if run in a multi-core system. In the snakemake command, specifying the number of cores (`--cores`) provides the information to snakemake to parallelize the tasks in the specific rules across the specified number of cores, thus utilizing multiprocessing. Within each rule, the tools being used also support inherent multithreading (for example, ASTRAL-Pro2, PASTA, MashTree, and FastTree support multithreading). Snakemake parallelizes the tasks across a specific number of cores provided by the user (`--cores`) in the main snakemake command. ROADIES performs multithreading by using `--threads` settings for individual snakemake rules and specifying the number of threads (`--num_threads`) while executing the snakemake command. If `--cores` are set to 32, each snakemake rule will be parallelized as 32 parallel instances across 32 cores (by default, one task consumes one thread or one core). If `--threads` is set to 4 for a rule, snakemake will parallelize that rule 8 times, each task consuming four threads or four cores, thus achieving a mix of multithreading and multiprocessing.

In ROADIES, we set the number of threads for sampling, LASTZ, PASTA, RAxML-NG, and MashTree rules across all modes of operation as the *number of cores/4* (four parallel instances will run for each rule). The number of instances allowed to run in parallel for each rule can be configured using the NUM_INSTANCES parameter in `config.yaml`. Each task run by PASTA, RAxML-NG, and MashTree is multithreaded to the number of threads equal to *cores/NUM_INSTANCES*. To avoid the memory bandwidth bottleneck and achieve efficiency due to parallelism, ROADIES has NUM_INSTANCES set to 4 by default. More parallel instances result in more memory bandwidth requirements (for larger genomes, it can lead to memory bottleneck). We would have more multithreading for the same task with fewer parallel instances. However, PASTA and RAxML-NG are proven to run optimally with four threads. Hence, setting the number of instances (NUM_INSTANCES) equal to the number of cores/4 is optimal. ASTRAL-Pro2 is also parallelized across the number of specified cores (`--cores`) given using the snakemake command.

### ROADIES: Configuration and Execution

The execution of the ROADIES pipeline involved three major steps. First, we downloaded and gathered the dataset containing all input genomic sequences in `.fa` format or `.fa.gz` format. The file names of genomic sequences corresponded to the respective species’ names (for example, the file containing genomic data for species Aardvark is named `Aardvark.fa`). Second, before running the pipeline, we configured the parameters in the YAML file (`config.yaml` - provided in the GitHub repository). We specified the input genomic dataset path (GENOMES) and the path where results will be saved (OUT_DIR) in the YAML file, along with some additional parameters crucial for executing ROADIES (detailed significance of each parameter along with its default values are mentioned in https://turakhia.ucsd.edu/ROADIES/). Third, after configuring the parameters, we run the wrapper script `run_roadies.py` using the command:

python run_roadies.py --cores {number of cores} --config {config file path} ---output {output directory} --converge (for running in converge mode --mode {mode of operation - accurate/fast/balanced}.

This script calls the underlying snakemake workflow described in previous sections, with the specified number of cores (`--cores`), the configuration file (`--config`; this is `config.yaml` by default), and the mode of operation (`--mode accurate` by default). Once the execution was completed, ROADIES generated the final species tree in the Newick format as `roadies.nwk` in the specified folder (specified as OUT_DIR parameter in the configuration file). The final species tree provided by ROADIES was an unrooted tree with branch lengths for all nodes of the tree.

### Execution environment and Runtime estimates

ROADIES pipeline was executed within a custom conda environment where all the required tools were installed (such as LASTZ, MAFFT, RAxML-NG, FastTree, and their prerequisites) used in various stages of the pipeline (`mamba create -y -c conda-forge -c bioconda --name roadies_env snakemake alive-progress biopython numpy lastz mashtree matplotlib seaborn treeswift=1.1.28 fasttree=2.1.11 python=3.11 raxml- ng ete3`). After creating and activating the conda environment (`conda activate roadies_env`), we installed ASTRAL-Pro2 and PASTA separately and built the C++-based sampling script (used in the first stage of the ROADIES pipeline). We prepared an automated executable script (`roadies_env.sh`) to download all tools and dependencies at once and activate the conda environment required to run the ROADIES pipeline.

We performed all the experiments discussed in the manuscript on AWS EC2 R6a instances (r6a.4xlarge), which is a 16-core memory-optimized instance suitable for memory-intensive workloads. The input datasets were downloaded from the NCBI Genome Browser (https://www.ncbi.nlm.nih.gov/gdv/) using the NCBI IDs listed in the Supplementary Tables 1-3 and saved in AWS elastic block storage (EBS) volumes in `.fa.gz` format, which were then provided as GENOMES parameter in the configuration file before running ROADIES.

To parallelize the ROADIES execution for higher GENE_COUNT value, we ran the ROADIES pipeline across multiple AWS instances simultaneously. For example, for GENE_COUNT=64000, the ROADIES pipeline was executed simultaneously in 16 different AWS EC2 instances (each instance of r6a.4xlarge) with GENE_COUNT=4000 in each AWS instance. Thus, we were able to parallelize ROADIES for 64000 genes in 16 separate blocks, each with 4000 genes. After the completion, we concatenated the gene tree list (`gene_tree_merged.nwk`) and mapping file (`mapping.txt`) from all parallel instances to get a master list of gene trees and mapping file corresponding to 64000 genes sampled across all AWS instances, required by ASTRAL-Pro2 to generate the final species tree (`roadies.nwk`).

At the end, ROADIES also outputs the estimated runtime in the `time_stamps.csv` file in seconds format. This runtime is the wall clock time consumed for executing all stages of the ROADIES pipeline from start to end. For the converge mode, ROADIES estimates the runtime for all the iterations in the same file, which is reported for all the experiments discussed in the manuscript.

### Cophylogenetic plots and accuracy estimates

ROADIES does not generate rooted trees by default. Hence, for easy comparison of the final species tree with the reference tree (which is provided through the REFERENCE parameter in the `config.yaml` file), we rerooted the final species tree generated by ROADIES such that its root position matches with the root of the reference tree. We provide a `reroot.py` script for the user to reroot the trees explicitly by providing a reference tree.

The species-level phylogenetic tree discussed in the manuscript is the final `roadies.nwk` output from ROADIES, from which an order-level phylogeny was semi-manually derived. After generating the species and order level tree for both ROADIES and the reference, we constructed the cophylogenetic plot to compare the tip displacements and topological differences between the trees when placed adjacently. All the phylogenetic tree illustrations are made using iTOL software^100^. We manually rotated the tips in the trees for better one-on-one mapping between both trees with minimal tip displacements.

ROADIES quantifies the difference between two trees using the normalized Robinson-Foulds distance as the distance metric. We used the ETE3 toolkit ^101^ to compare two trees, `tree1` and `tree2`, using the function `tree2.compare(tree1)` to estimate normRF.

### Stability and scalability analysis

The stability and scalability tests are performed on AWS EC2 instances. To perform the stability test, we perform ROADIES pipeline execution with 100 drosophilid genomes and the same set of configuration parameters (GENE_COUNT=4000) on four 16-core AWS EC2 instances simultaneously. After getting the final species trees from 4 independent instances, we calculated the normRF for all four species trees with the reference tree. Following the same approach, we performed the scalability test, with the difference of providing different numbers of genomes and varying GENE_COUNT as input across four AWS instances.

### Baseline methods

ROADIES follows an automated approach to estimating phylogenetic trees from raw genomic sequences. In contrast, state-of-the-art methods involve computationally expensive steps, including gene annotations, whole genome alignments, orthology inference, and multiple sequence alignment. To compare the performance of ROADIES, we wanted to compare it with the conventional pipeline replicating all the stages executed by ROADIES. However, it is difficult to replicate a traditional pipeline completely. Hence, to estimate the runtime of the baseline and compare it with ROADIES, we only accounted for whole genome alignments and orthology inference, which is performed by TOGA ^39^.

For the comparative analysis of ROADIES with TOGA, we executed the TOGA pipeline between a pair of mammalian genomic sequences (genomic sequence of human - hg38 assembly (*Homo sapiens*) and mouse - mm10 assembly (*Mus musculus*)) in a 16-core AWS EC2 instance (https://github.com/hillerlab/TOGA). However, for executing TOGA, chain-formatted alignments are essentially required as the input, for which we collected the human and mouse assemblies in

`.fa` format and performed the LASTZ chaining (https://github.com/hillerlab/make_lastz_chains) using the command: `/make_chains.py target query hg38.fa mm10.fa --pd test_out -f --chaining_memory 16` to get the output `hg38.mm10.final.chain.gz`. Next, we collected the `.isoforms.tsv`, `.U12sites.tsv`, and `.bed` files from the TOGA repository provided for the hg38 assembly, along with the genomic sequences of reference and query, and ran TOGA using the command: `./toga.py`

../make_lastz_chains/test_out/hg38.mm10.final.chain supply/hg38.v35.for_toga.bed hg38.2bit mm10.2bit --kt --pn test -i

`supply/hg38.v35.for_toga.isoforms.tsv --cjn 16 --u12 supply/hg38.U12sites.tsv --ms`. To estimate the runtime for both LASTZ chaining and TOGA, we used the wall clock time using the `time` command in the terminal.

Considering ROADIES follow a fully automated approach, MashTree ^24^ is the only tool we are aware of that falls under the same automated category of inferring species trees directly from raw genomic sequences. Hence, to compare the performance of ROADIES with MashTree, we installed MashTree in our system of 16-core AWS EC2 instance (r6a.4xlarge) (https://github.com/lskatz/mashtree) and generated the species tree by running the following command: `mashtree --mindepth 0 --numcpus 16 *.fastq.gz [*.fasta] > mashtree.dnd`. We used the `time` command to estimate the runtime and recorded the wall clock time in the manuscript. From the final MashTree output, we constructed the order-level/group-level cophylogenetic tree in the same way mentioned in the section “cophylogenetic plots and accuracy estimates.”

## Acknowledgments

We thank Tian (Kevin) Liu for his valuable contributions to the ROADIES codebase. We thank Hiram Clawson, Guojie Zhang, Yulong Xie, and Benedict Paten for their helpful feedback. Research reported in this publication was supported by an Amazon Research Award (Fall 2022 CFP), NIH grant 1R35GM142725 (to S.M.), and funding from the Hellman Fellowship (to Y.T.).

## Supplementary Figures

**Supplementary Figure 1:**
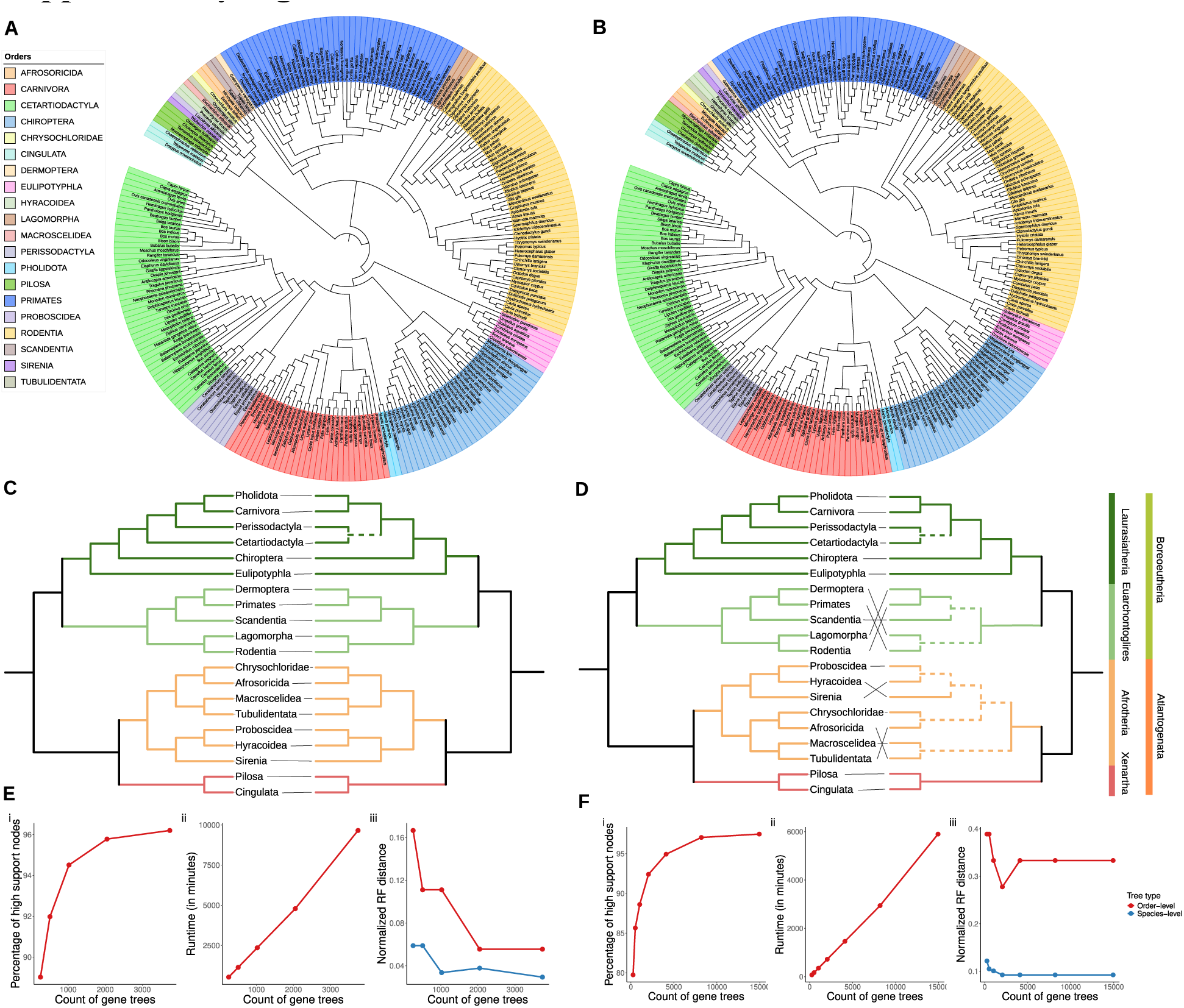
ROADIES results evaluated on 240 placental mammals (in the balanced and fast modes of operation). **(A-B)** The species-level phylogenetic tree of 240 placental mammals estimated by ROADIES in (A) the balanced mode and (B) the fast mode**. (C- D)** Order-level cophylogenetic tree of 240 placental mammals estimated by ROADIES in (C) the balanced mode and (D) the fast mode compared to the reference tree by Zoonomia consortium ^12^. **(E-F)** ROADIES convergence results in (E) the balanced mode and (F) the fast mode. As the number of gene trees increases, we show the percentage of highly supported species tree nodes with localPP >= 0.95 (plot i), the linear increase in runtime (ii), and the normalized Robinson Foulds distance of the final species tree to the reference tree (iii).

**Supplementary Figure 2:**
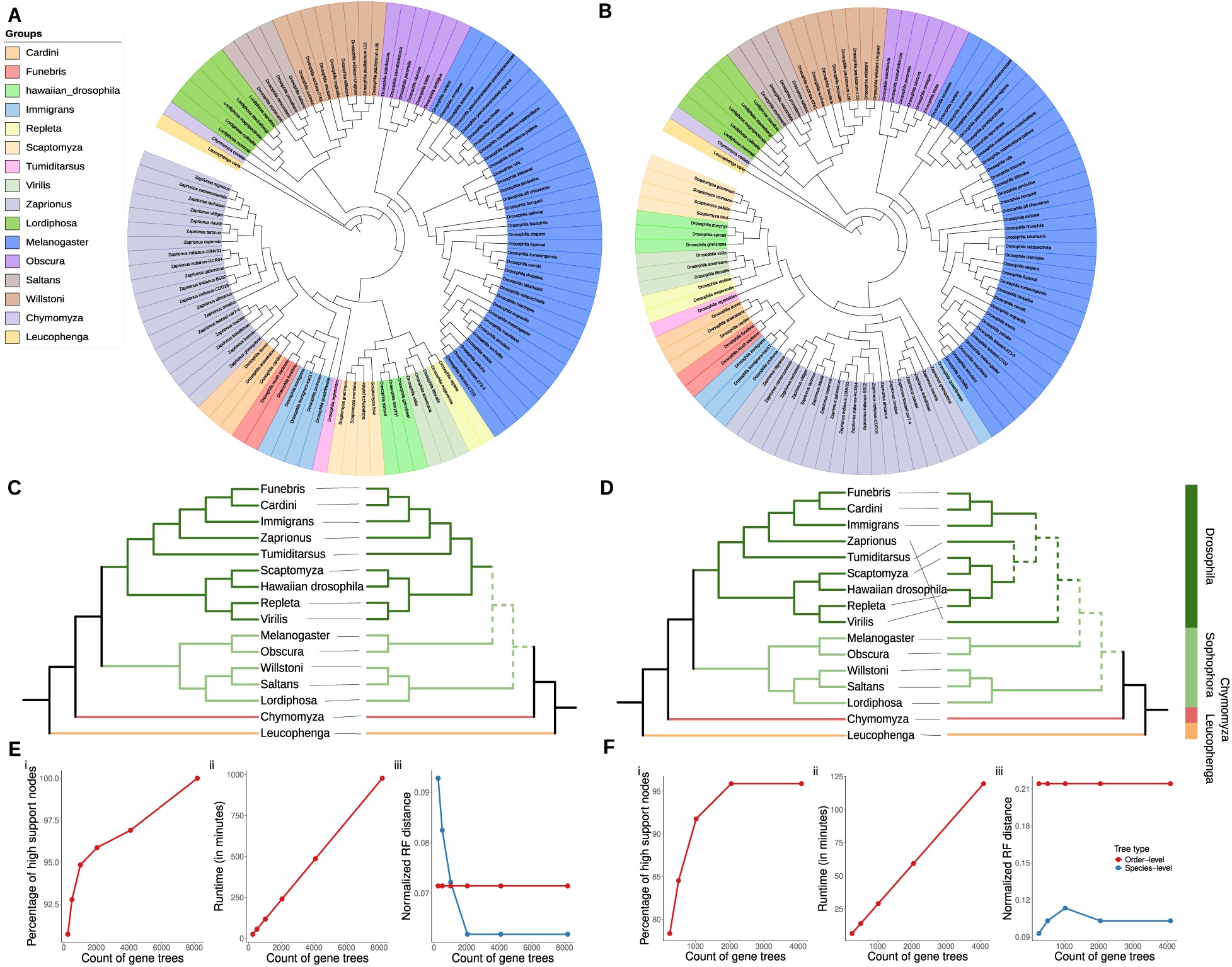
ROADIES results evaluated on 100 drosophilid species (in the balanced and fast modes of operation). **(A-B)** The species-level phylogenetic tree of 100 drosophilid species estimated by ROADIES in (A) the balanced mode and (B) the fast mode**. (C- D)** Group-level cophylogenetic tree of 100 drosophilid species estimated by ROADIES in (C) the balanced mode and (D) the fast mode compared to the reference tree by Kim et al. ^30^. **(E-F)** ROADIES convergence results in (E) the balanced mode and (F) the fast mode. As the number of gene trees increases, we show the percentage of highly supported species tree nodes with localPP >= 0.95 (plot i), the linear increase in runtime (ii), and the normalized Robinson Foulds distance of the final species tree to the reference tree (iii).

**Supplementary Figure 3:**
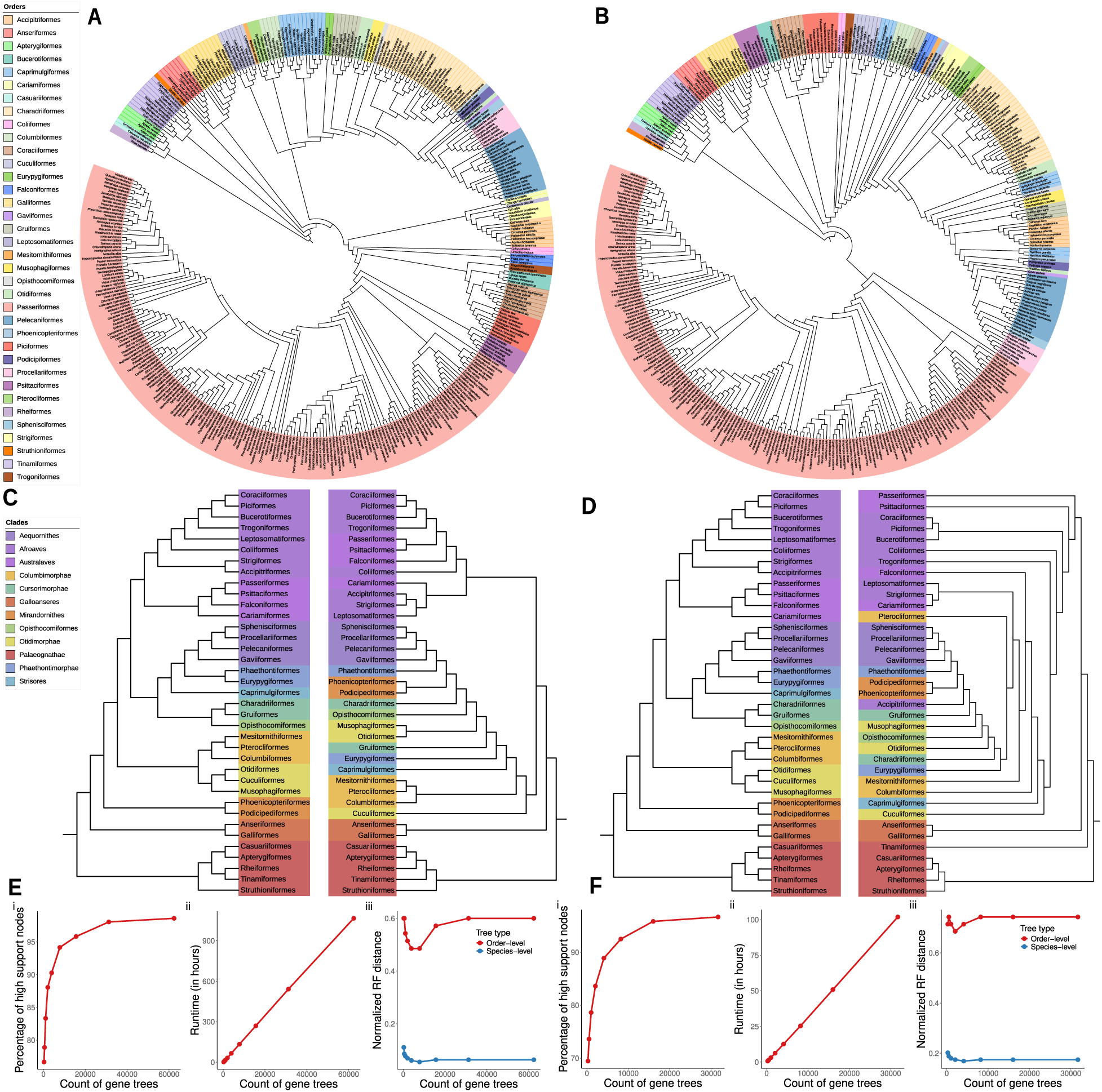
ROADIES results evaluated on 363 avian species (in the balanced and fast modes of operation). **(A-B)** The species-level phylogenetic tree of 363 avian species estimated by ROADIES in (A) the balanced mode and (B) the fast mode**. (C-D)** Order-level cophylogenetic tree of 363 avian species estimated by ROADIES in (C) the balanced mode and (D) the fast mode compared to the reference tree by Stiller et al. ^6^. **(E-F)** ROADIES convergence results in (E) the balanced mode and (F) the fast mode. As the number of gene trees increases, we show the percentage of highly supported species tree nodes with localPP >= 0.95 (plot i), the linear increase in runtime (ii), and the normalized Robinson Foulds distance of the final species tree to the reference tree (iii).

**Supplementary Figure 4:**
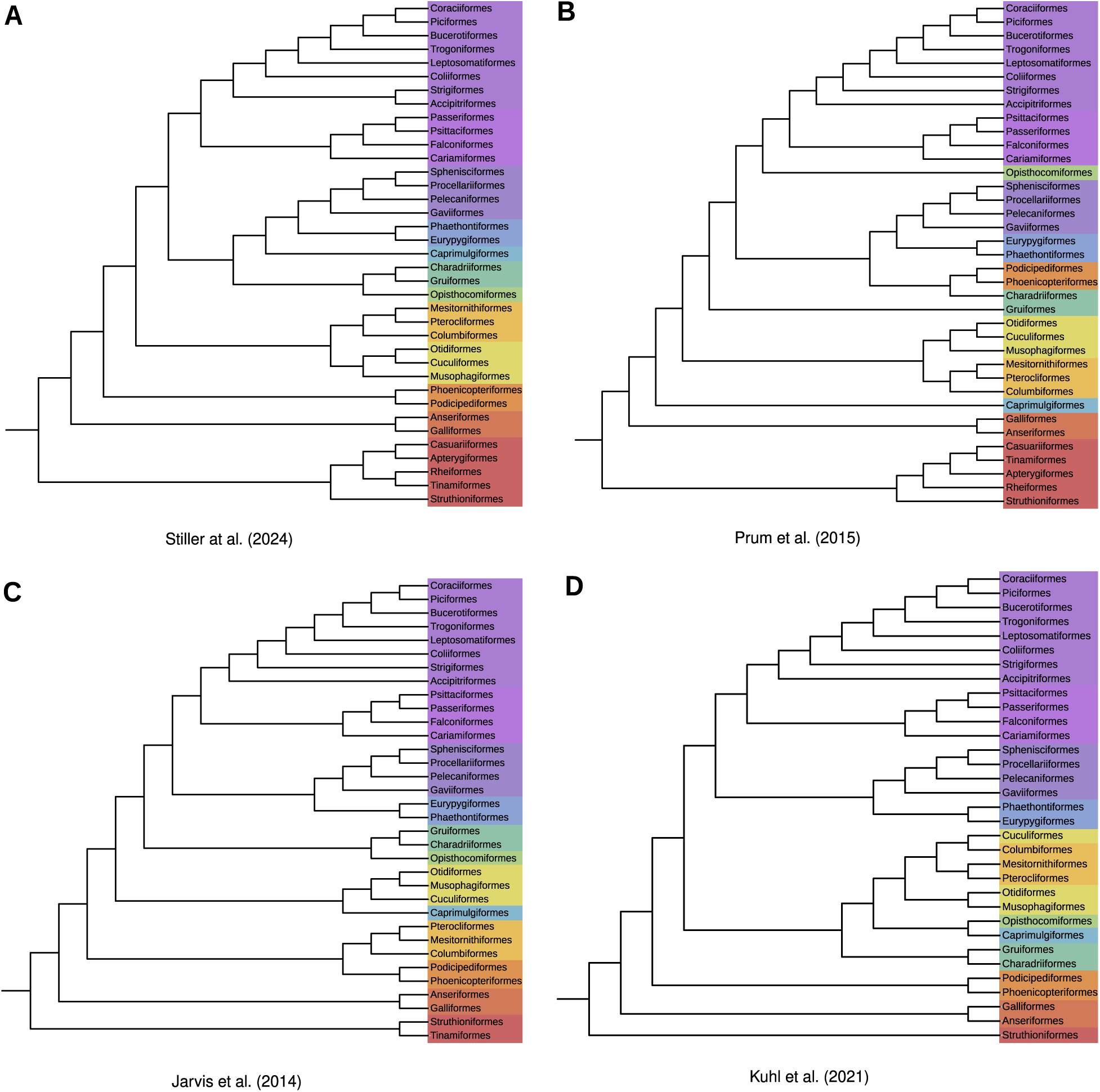
Comparison of Birds phylogeny at the order level from different established approaches. **(A)** The order-level phylogenetic tree of 363 avian species proposed by Stiller et al. ^6^ using 63,430 loci (44,846 intronic, 14,972 exonic, and 4,985 UCE loci). This tree is the golden reference for our entire analysis. **(B)** Order-level phylogenetic tree of 198 avian species proposed by Prum et al. ^68^ **(C)** Order-level phylogenetic tree of 48 birds proposed by Jarvis et al. ^4^ **(D)** Order-level phylogenetic tree of avian species proposed by Kuhl et al. ^67^ by analyzing 3’- UTRs of 221 avian family-level taxa including 379 genera and 429 species.

**Supplementary Figure 5:**
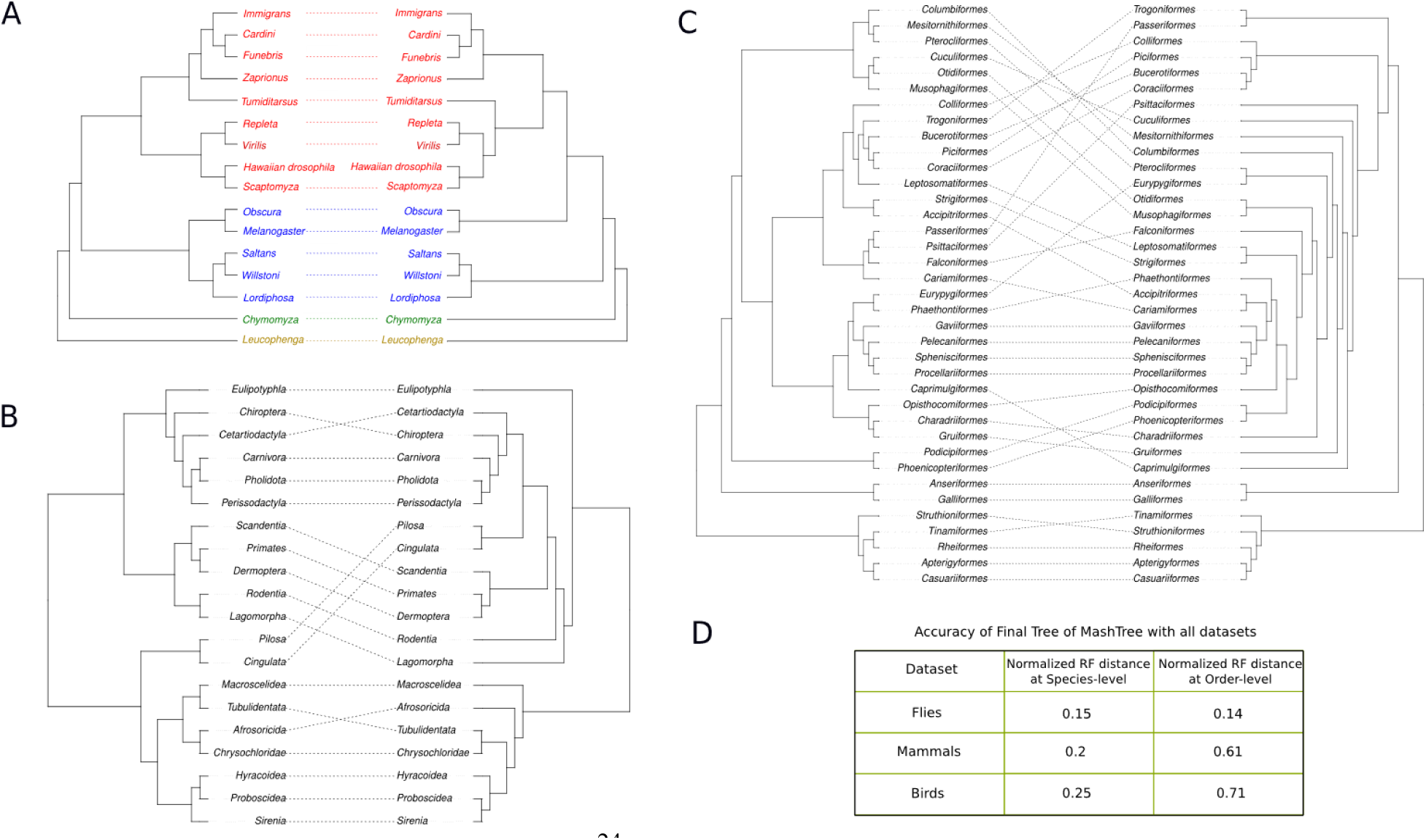
MashTree^24^ results evaluated on all datasets. **(A)** Group-level cophylogenetic tree of 100 drosophilid species estimated by MashTree (right) compared to the reference tree from Kim et al. ^30^ (left). **(B)** Order-level cophylogenetic tree of 240 placental mammals estimated by MashTree (right) compared to the reference tree from Zoonomia consortium ^12^ (left). **(C)** The order-level cophylogenetic tree of 363 avian species estimated by MashTree (right) compared to the reference tree from Stiller et al. ^6^ (left). **(D)** Normalized Robinson-Foulds distance (at the order-level/group-level and species-level) between the reference tree and the tree inferred by MashTree for all three sets of genomes.

**Supplementary Figure 6:**
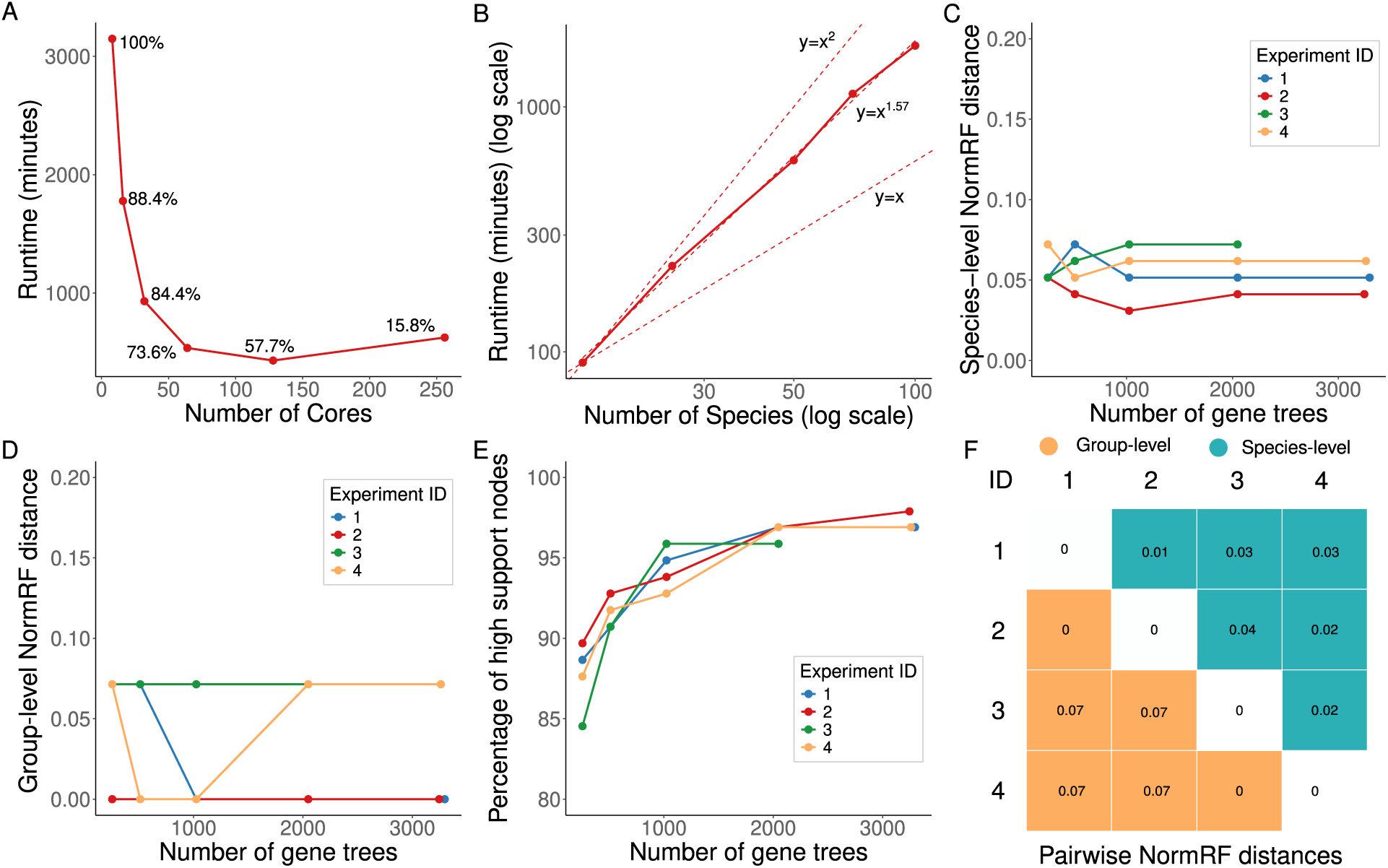
ROADIES scaling and variance results with 100 Drosophilid species in accurate mode. **(A)** The runtime of the entire ROADIES pipeline, with increasing system cores and the same set of parameters (GENE_COUNT = 4000), exhibiting strong scaling. Data points show the efficiency compared to the baseline of ROADIES runtime with 8 cores. **(B)** The runtime of the ROADIES pipeline with an increasing number of species, all in 16 core systems, and with GENE_COUNT = 4000. ROADIES scales with a slope of 1.57 (between quadratic and linear) with increasing species count. **(C-E)** Variance of results by running multiple ROADIES experiments in 16 core machines, all with GENE_COUNT = 4000. **(F)** Pairwise NormRF distances by comparing ROADIES output from a unique pair of experiment IDs.

**Supplementary Figure 7:**
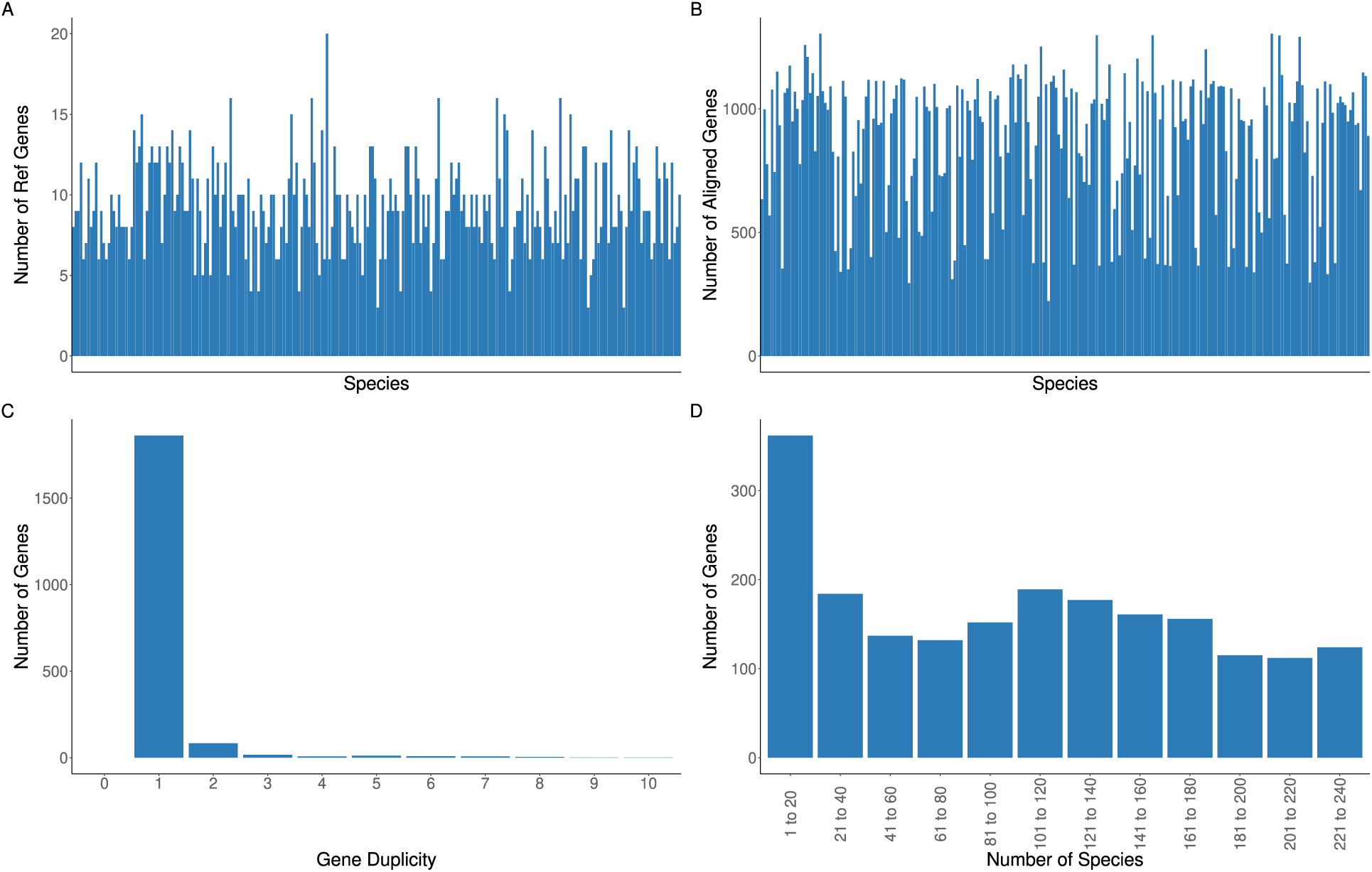
Additional statistical plots generated by ROADIES for 240 placental mammals in accurate mode. **(A)** ROADIES results with random sampling for 240 Placental Mammalian species in accurate mode. The plot shows the number of genes sampled (Y-axis) from each genome (X-axis) across the entire input dataset, with total GENE_COUNT = 2000. **(B)** ROADIES results with pairwise alignment for 100 Drosophilid species in accurate mode. The plot shows the number of genes (Y-axis) aligned to each genome (X-axis) across the input dataset, with GENE_COUNT = 2000. **(C)** Histogram plot of the count of the genes (Y-axis) and the number of their corresponding duplicate copies in the same species (X-axis). (GENE_COUNT = 2000). **(D)** Histogram plot of the count of genes (Y-axis) spanning the specified number of homologs (X-axis).

**Supplementary Figure 8:**
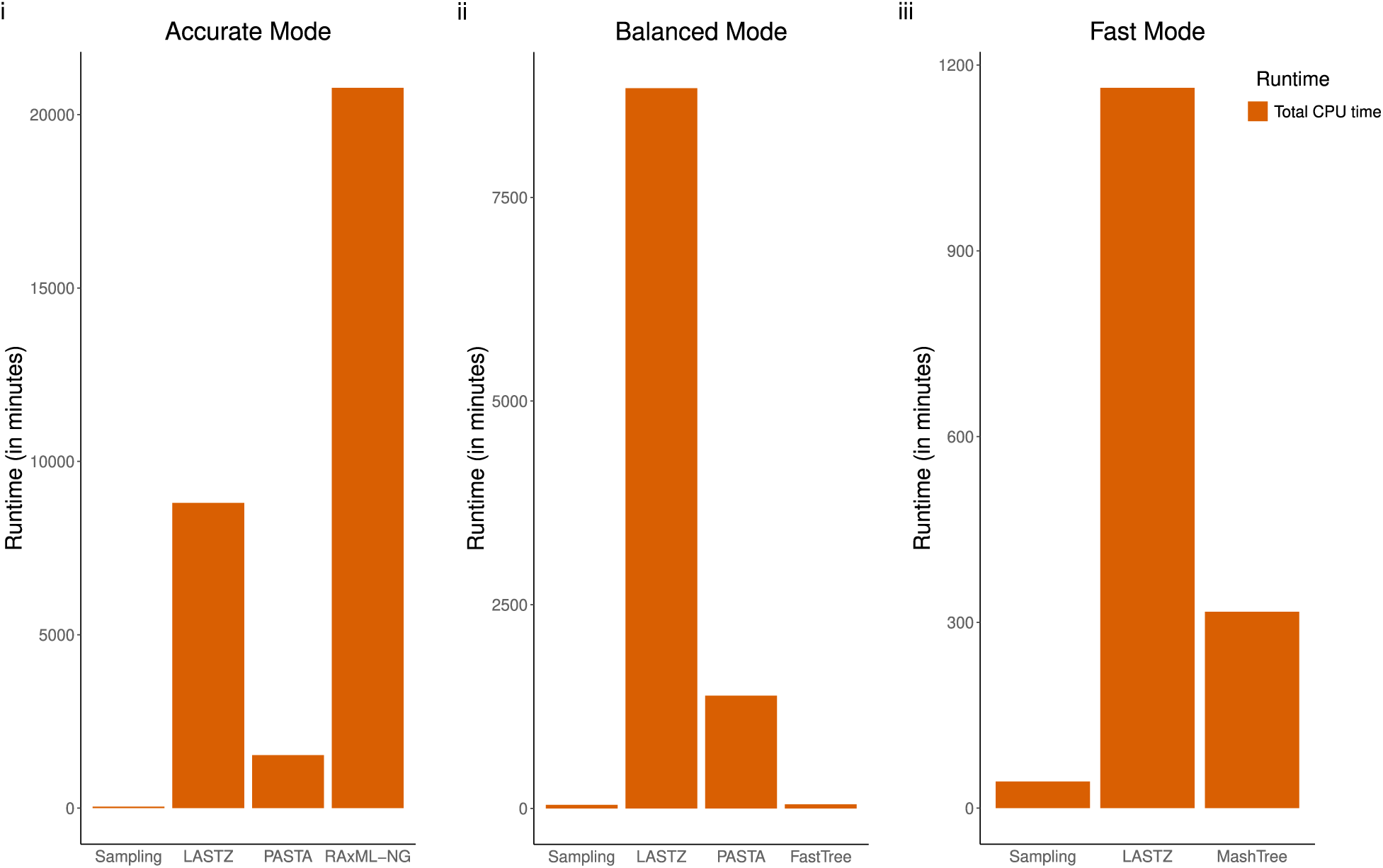
Stagewise CPU time estimate for all modes of operation (with 240 placental mammals, GENE_COUNT = 2000). All experiments were performed on the 16-core AWS R6a instance, consuming a total wall clock time of around 113h, 81h18m, and 12h8m respectively.

